# Physical and data structure of 3D genome

**DOI:** 10.1101/596262

**Authors:** Kai Huang, Yue Li, Anne R. Shim, Rikkert J. Nap, Vasundhara Agrawal, Ranya K.A. Virk, Adam Eshein, Luay M. Almassalha, Vadim Backman, Igal Szleifer

## Abstract

With the textbook view of chromatin folding based on the 30nm fiber being challenged, it has been proposed that interphase DNA has an irregular 10nm nucleosome polymer structure whose folding philosophy is unknown. Nevertheless, experimental advances suggested that such irregular packing is associated with many nontrivial physical properties that are puzzling from a polymer physics point of view. Here, we show that the reconciliation of these exotic properties necessitates modularizing 3D genome into tree data structures on top of, and in striking contrast to the linear topology of DNA double helix. Such functional modules need to be connected and isolated by an open backbone that results in porous and heterogeneous packing in a quasi-self-similar manner as revealed by our electron and optical imaging. Our multi-scale theoretical and experimental results suggest the existence of higher-order universal folding principles for a disordered chromatin fiber to avoid entanglement and fulfill its biological functions.

## Introduction

The intimate connection between 3D interphase DNA structure and gene expression in eukaryotic cells has made chromatin folding a rapidly developing field. In the last decade, previously well-accepted concepts have been continuously challenged by new experimental discoveries. For example, it had previously been widely believed that chromatin is folded into 30nm fibers^1^, which are assembled into discrete higher-order structures. However, such regular folding hierarchy has not been observed in the state-of-the-art electron microscopy tomography (ChromEMT) experiment^2^, which instead revealed a highly disordered chromatin polymer that is heterogeneously packed even down to the level of single nucleosomes in situ. This structural heterogeneity of chromatin has been suggested by Partial Wave Spectroscopy (PWS)^3^, a label-free technique, to have a profound regulatory impact on the global transcriptional profile of live single cells^4^. At the population-average level, thanks to the development of chromosome conformation capture (3C)^5^ and related techniques, it is becoming well established that chromatin has frequent genomic contacts inside topologically associating domains (TADs)^6^. A loop extrusion hypothesis based on CTCF and cohesin proteins was proposed to explain many TAD features at the ensemble level^7,8^. However, the importance of this putative mechanism at the single-cell level has been questioned by recent super-resolution fluorescence microscopy experiment^9^, which found chromatin segments to be rich in TAD-like clusters^10^ that are persistent even after cohesin knockout. Taken together, these new experimental findings remind us of the complexity of chromatin folding, which requires the cooperation of a large family of architectural chromatin-regulatory proteins, as well as the interplay between multiple folding mechanisms such as supercoiling^11–14^, phase separation^15–17^, molecular binding^18^, crowding effects^19^ and loop extrusion^7^, all under the feedback control of transcription to be responsive to external stimuli. They also raise many important questions such as (1) what are the functional units of chromatin, (2) what is the hierarchy of chromatin folding hidden in the disordered morphology, and (3) what are the inner workings of TADs at the single-cell level. Among these questions, perhaps the most fundamental one is whether there are abstract yet universal folding principles of our genomic code independent of the known molecular and mechanistic complexity.

To investigate the existence of such principles (mathematical rules) without the burden of accounting for all the physical interactions and biological mechanisms that are far from fully understood, one could search for an abstract folding algorithm that aims to recapitulate the major experimental observations with minimal adjustable parameters and computational complexity, the success of which would suggest a positive answer. Among the chromatin features known to date from experimental results, the most important two are (1) the frequent self-interactions^6^ that link promoters and enhancers for transcriptional regulation, and (2) the heterogeneous packing^2,20^ that disperses local DNA accessibilities, making room for transcription and nuclear transport. However, there is an apparent conflict between these two major chromatin properties from a polymer physics point of view. For example, it has been hypothesized that chromatin resembles a fractal globule (FG)^21,22^, which is a self-similar polymer in a fully collapsed state. While the FG model predicts the observed high contact frequency, it cannot explain the spatial heterogeneity of chromatin packing. In fact, one can prove that it is impossible for any 1D chain to be simultaneous self-similar, rich in self-contacts, and diverse in packing density^23^ (see supplementary materials for more details). This trilemma rules out the possibility of chromatin being any kind of fractal chain (Fig. 1A, Fig. S1) and means that chromatin must have distinct folding modes at different length scales. Despite the unknown folding details, such scale dependence has manifested itself in the mass scaling measured by small angle neutron scattering^24^, which undergoes a transition from slow to fast as the length scale increases (Fig. 1B). Since slow mass scaling is associated with polymer decondensation and therefore less intra-polymer contacts, one would expect chromatin contact frequency to decay faster at a smaller scale. However, high throughput 3C (Hi-C) experiments indicate a contact scaling transition from slow to fast as the genomic scale increases (Fig. 1C), opposite to the expected result from mass scaling. Moreover, at the small length scale where mass scaling is not even close to 3 (i.e., space filling), chromatin exhibits a remarkably slow power-law decay of contact frequency with a scaling exponent around - 0.75 inside TADs^7^. Such abnormal behaviors make chromatin folding not only an intriguing biological question but also a fascinating puzzle for polymer physics.

**Figure 1.**
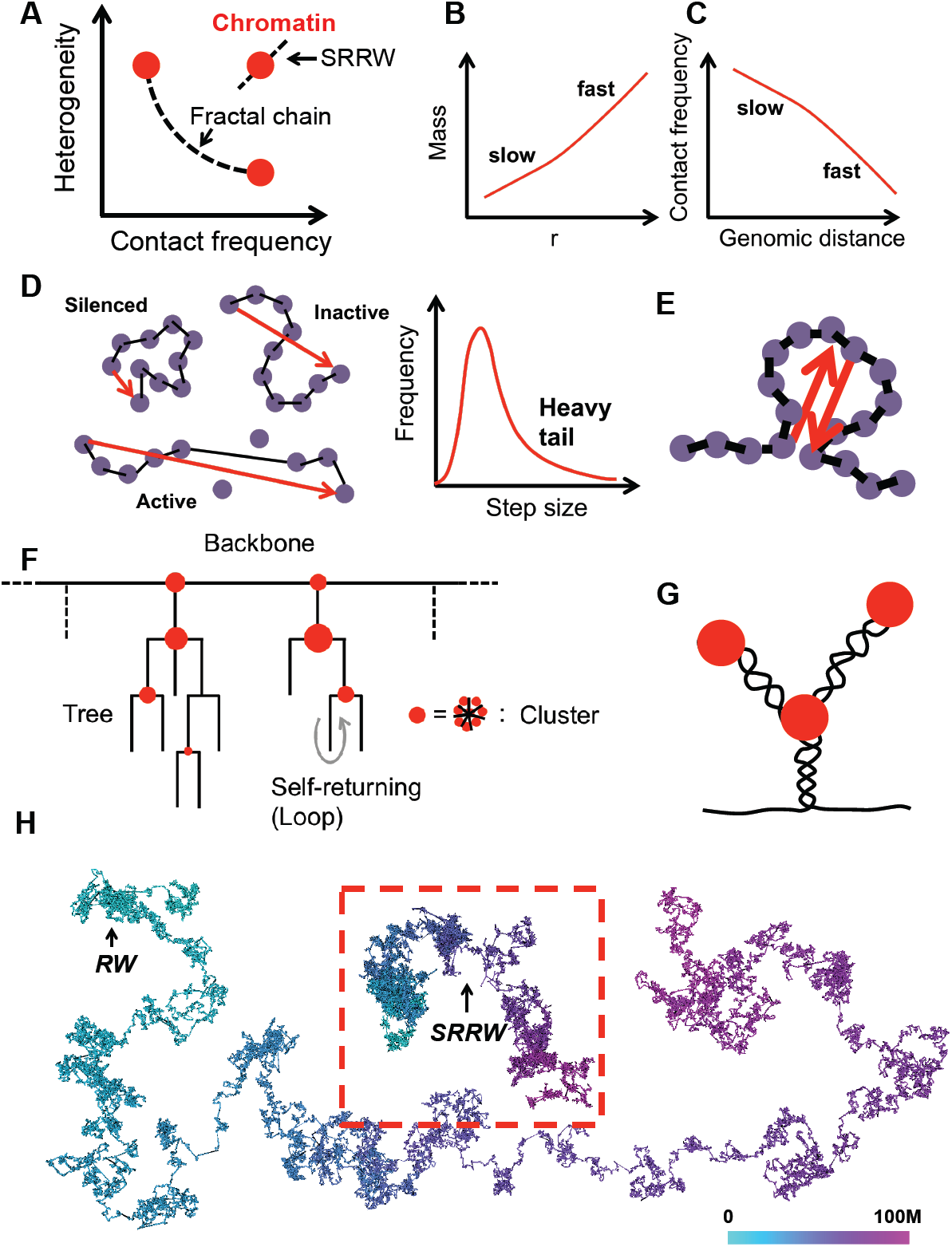
The paradox of chromatin folding and the basic ideas of SRRW. (**A**) Chromatin structure cannot be explained by a fractal chain that has anti-correlated contact frequency and density heterogeneity. SRRW is developed to unify these two properties. (**B**) Schematic representation of the mass scaling behavior of chromatin. (**C**) Schematic representation of the contact scaling behavior of chromatin (<10Mb), which is counterintuitive given the mass scaling behavior. (**D**) Coarse-grained representation of diverse epigenetic states at nano-scale using steps with a wide distribution of step sizes. One step approximately maps to 10 nucleosomes or 2kb DNA. (**E**) Self-returning as a coarse-grained representation of looping (in a broad sense including supercoiling and clustering). (**F**) SRRW’s topological architecture featuring random trees connected by a backbone. Tree nodes are formed by frequent self-returning of short steps. (**G**) One possible realization of tree structures is by combining nanoclusters with nested supercoils or loops. (**H**) Comparison between a free SRRW and a free RW, both of 50,000 steps (100Mb)

In this paper, we address the above contact-structure paradox by introducing a self-returning random walk (SRRW) as a mathematical model that effectively breaks the non-branching topology of the 10nm chromatin fiber and generates tree-like topological domains connected by an open chromatin backbone. The decondensed backbone segments isolate the topological domains and allow them to form larger compartments with high folding variation. We carry out nanoscopic imaging using chromatin scanning transmission electron microscopy (ChromSTEM) and live-cell Partial Wave Spectroscopic (PWS) microscopy to demonstrate DNA packing heterogeneity across different scales, visualizing chromatin domains and compartments at the single-cell level as predicted by our model. Our results support the hypothesis of local DNA density being an important transcriptional regulator and provide a new picture of genomic organization in which chromatin is folded into a variety of minimally-entangled hierarchical organizations across length-scales ranging between tens of nanometers to microns, without requiring a 30nm fiber. We discuss the quantitative predictions of this folding picture and how they explain a wide range of existing experimental observations. Using heat shock as a model system, we further reveal couplings between chromatin properties during stress response. The strong agreement between our model and experiments on this structural perturbation sheds lights into the structure-function relationship of interphase DNA and suggests the existence of higher-order folding principles and significant reduction of dimension during genomic landscape exploration. We end the paper by discussing possible molecular mechanisms and biological consequences of this novel organizational paradigm.

## Self-returning random walk (SRRW)

Given the enormous size of our genome, coarse-graining is necessary in chromatin modeling. Typically, polymer models coarse-grain at least a few kb of DNA into one monomer bead in order to study chromatin structures above 10Mb. This implicitly restricts the basic repeated unit of the coarse-grained chromatin to be of 30nm or larger in diameter^25^, incompatible with the recent ChromEMT observation where chromatin is predominantly a heterogeneous assembly of 5 to 24nm fibers^2^. Therefore, instead of large beads of one size, we use steps with a continuous spectrum of step sizes to cover the conformational freedom^26–28^ of a 10nm fiber at the kb level (see Fig. 1D). We choose each single step to correspond with 2kb of DNA, roughly an average of 10 nucleosomes. To capture the frequent genomic contacts, we introduce stochastic, self-returning events (Fig. 1E). The return probability is assumed to decay with the length of the current step size *U*_0_ by a power law:

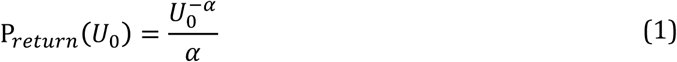

where *α* > 1 is the folding parameter for the modeled chromatin. Here, a smaller *α* leads to higher return frequency and vice versa. Once a step is returned, a further return is possible based on the same probability function. With a probability of 1-P_return_, a jump from the current position can be issued to explore a new point. The jump takes an isotropic direction and a step size *U*_1_ that follows a power law distribution:

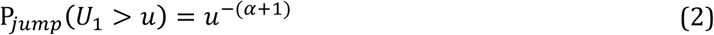

where *u* is larger than 1, with the resulting smallest step size in reduced units corresponding to roughly 30nm in the end-to-end distance of the 2kb nucleosome segment. Likewise, we eliminate unrealistically long step sizes, which correspond to 0.1 percent of the jumps. To model the confinement effect inside nucleus, we also introduce a global cutoff of 2*α* times the local cutoff during the conformation generation, which results in a DNA density of ∼0.015bp/nm^3^, comparable to that of an average diploid human eukaryotic nucleus. An accumulated stack of steps tracks the overall conformation of chromatin, whilst a subset of steps result in the formation of a chromatin “backbone”. By adding returning to jumping, the model thus turns a non-branching topology into a branching one with the degree of structural hierarchy controlled by the folding parameter *α*. As schematically shown in Fig. 1F, the overall topological architecture of SRRW is a string of random trees with the branches formed by the low-frequency returning of long steps and the nodes formed by the clustering of high-frequency returning of short steps. Isolated by the unreturned long backbone segments, the trees integrate nested loops and clusters into domains for co-regulation. One possible realization of such hierarchical structures can be the combination of passive nano-scale phase separation with active supercoiling driven by DNA transcription, as shown in Fig. 1G. Nested loops formed by molecular binding or extrusion could also contribute to the effective branching of chromatin. In the rest of the paper we use 50,000 steps to model 100Mb of DNA, roughly the average genomic size of one entire chromosome of human. We employ an *α* around 1.15 to generate structures that resembles interphase chromatin, and will discuss the implication of this parameter on higher-order chromatin folding.

## Chromatin structure and scaling at the single-cell level predicted by SRRW

At a negligible computational cost, SRRW is able to stochastically generate chromatin-like conformations at 2kb resolution of high case-to-case variations in spatial organization, but with consistent topological and statistical characteristics controlled by the global folding parameter *α*. Due to the hierarchical folding, a typical conformation generated by a free SRRW is much more compacted than that generated by a free random walk (RW) as shown in Fig. 1H. Since we are modeling one single interphase chromosome confined by the surrounding genome, we will focus on a confined SRRW (we keep the generic term SRRW for simplicity) as our chromatin model in the rest of the paper. As shown in Fig. 2A, the overall structure of our modeled chromatin is porous, non-globular^29^, with a rough surface, which is in stark contrast to the predictions of many polymer simulations where the modeled chromatins collapse into globules. The irregular shape of SRRW with large surface area would naturally facilitate accessibility of DNA to transcription inside interchromosomal domains between chromosome territories^30^, in line with experimental observations^31^. The rich porosity and sponge-like structure of SRRW also allow transcription factors to efficiently search the interior of chromatin and could explain the fractal-like nuclear diffusion of various biomolecules^32^.

**Figure 2.**
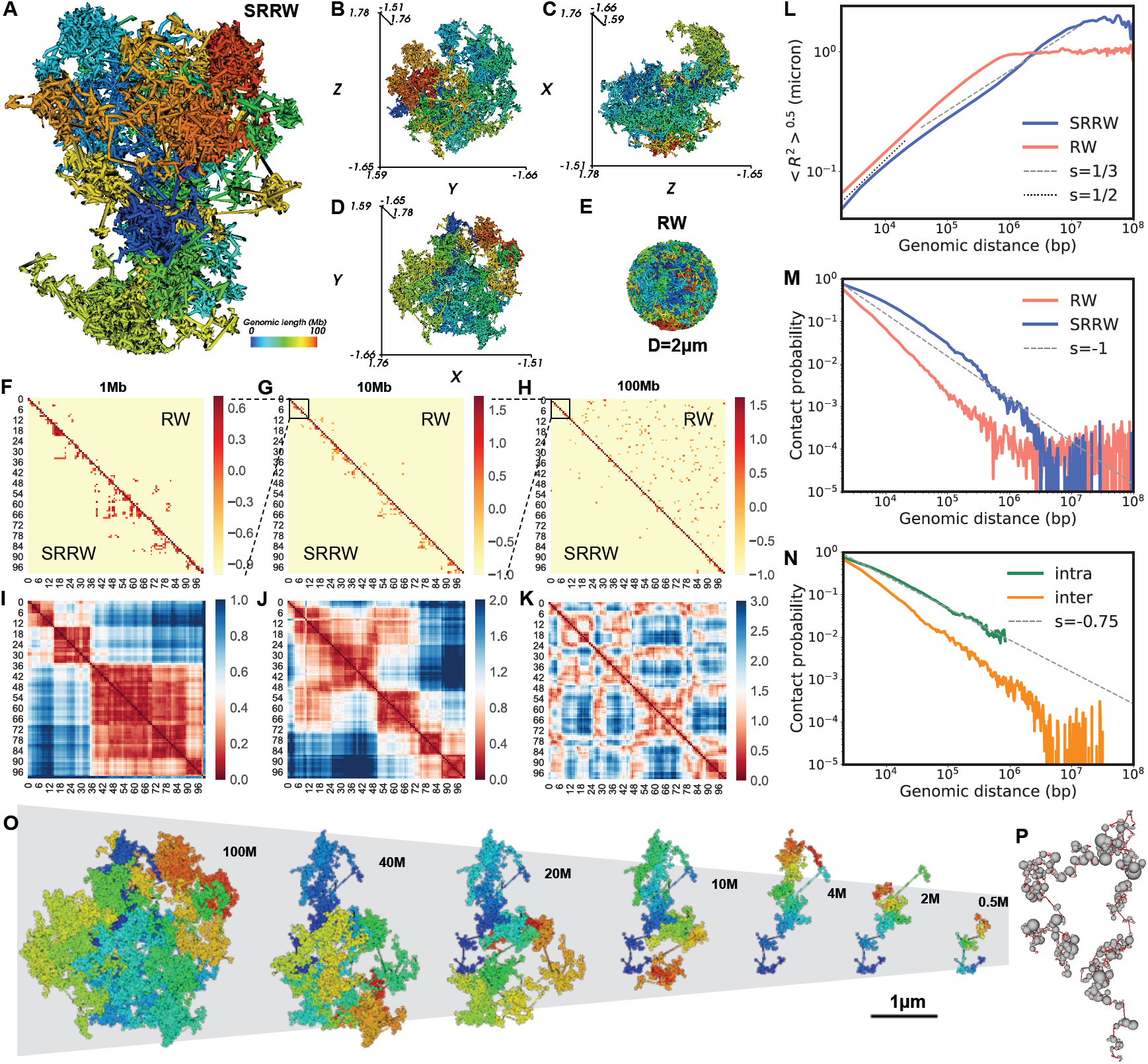
Typical single-cell level chromatin structure and its scaling behaviors predicted by our model. (**A-D**) Folded structure of our modeled chromatin and its xyz projections. (**E**) Equilibrium globule (confined RW) as a reference system. (**F-H**) Predicted single-cell level contact maps from local (1Mb) to global (100Mb) based on SRRW and RW. (**I-K**) The physical distance maps of SRRW from local to global. (**L**) Root-mean-squared end-to-end distance (*R*) scaling of SRRW and RW. (**M**) Contact scaling of SRRW and RW. (**N**) Contact scaling of SRRW for intra- and inter-tree contacts. (**O**) Structures of the modeled chromatin at different genomic scales. (**P**) Beads-on-string representation of the “secondary” structure of chromatin modeled by a free SRRW. Tree domains are represented in beads and backbone in string.

Colored based on the 1D genomic sequence, the modeled chromatin is clearly folded into domains and compartments reflected by the unmixed color. As a reference, an equilibrium globule (a confined RW, or RW for simplicity) shows no sign of territorial organization (Fig. 2E). The disparate organizations between SRRW and RW lead to strikingly different contact maps as displayed in Fig. 2F-H, where our modeled chromatin exhibits enriched contacts of TAD-like patterns^29,33,34^ in the most regulation-relevant ranges (≪10Mb) and is relatively depleted of random contacts above 10Mb. To better illustrate the spatial organization of the modeled chromatin, we calculated the physical distance matrices at different scales as shown in Fig. 2I-K. The TAD-like domains at the kb-to-Mb scale are clearly seen without requiring ensemble averaging, which is in consistence with recent super-resolution observations of heterogeneous domains in single cells^9^. To allow a more direct comparison between our predicted structures and the state-of-the-art imaging results^9^, we have adapted the experimental resolution of 30kb by coarse-graining our chromatin model 15-fold. A collection of typical structures of 2Mb modeled chromatin segments in both 2kb and 30kb resolutions and their physical distance maps are shown in Fig. 3, which demonstrates how 3D chromatin clustering leads to various TAD-like 2D patterns. More interestingly, our model predicts that such clustered contacts at the sub-Mb scale transition into plaid-like patterns at the multi-Mb scale (Fig. 2K), reminiscent of the AB-compartment patterns^5^ found in Hi-C contact maps. Our distance matrices imply that the 3D genome modeled at the single-cell level is compartmentalized across many scales, which is visualized in Fig. 2O. The compacted yet minimally intermixed 3D packing below 10Mb is reflected in the root-mean-squared end-to-end distance (*R*) scaling with slope between 1/3 and 1/2 in the corresponding regime (Fig. 2L). At the genomic scale above 10Mb, the slope drops to below 1/3, in line with experimental observation^35^ and signifying a strong intermixture of large compartments. Nevertheless, at this scale the modeled chromatin intermixes as clumps rather than wires, which would be expected to result in strong internal friction rather than entanglement.

**Figure 3.**
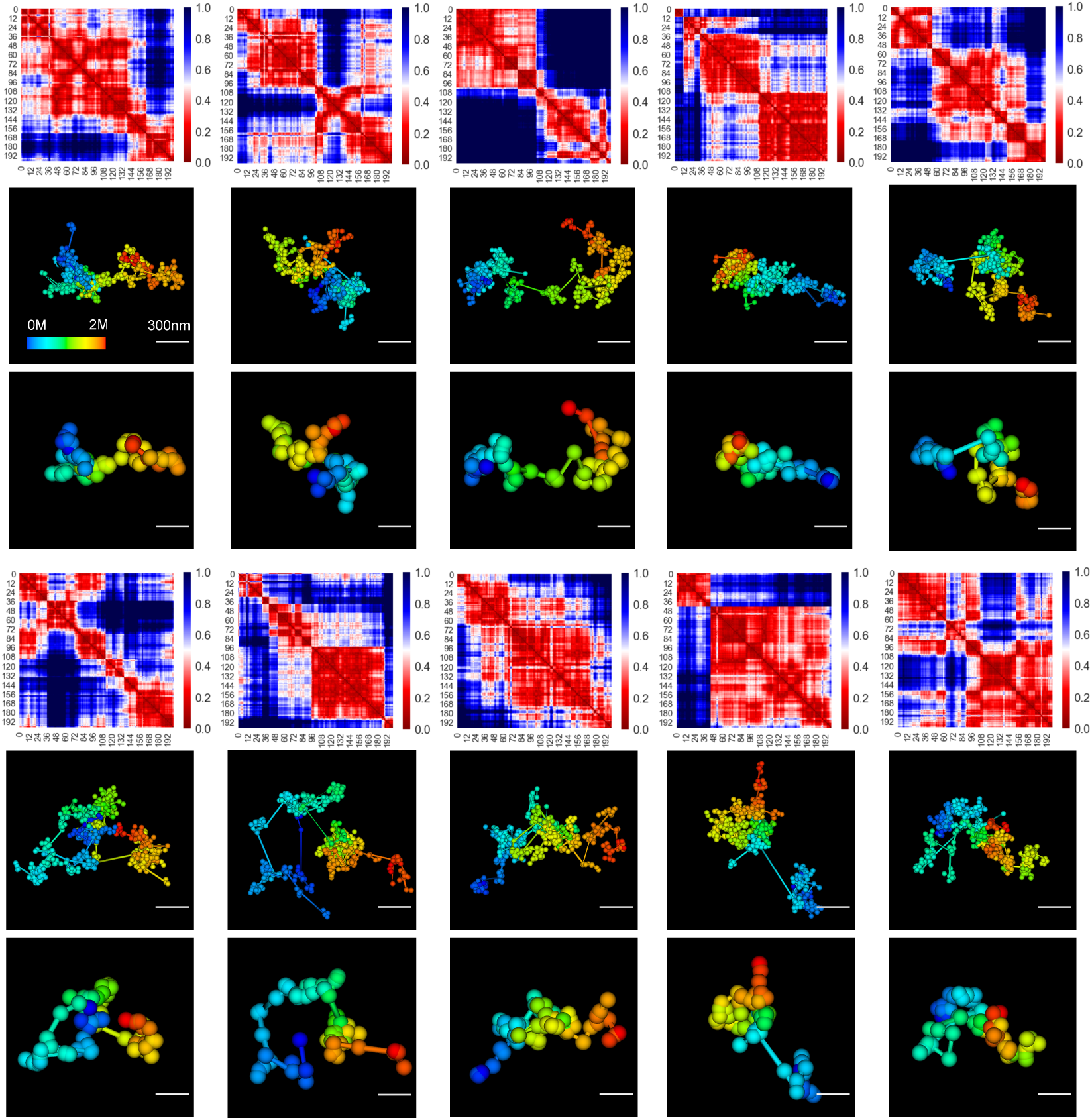
Typical chromatin structures of 2Mb of DNA at 2k and 30kb resolutions and their physical distance maps (in micron) at the single-cell level predicted by the SRRW model.

Fig. 2M shows the contact probability (*P*_*c*_) curve of our modeled chromatin. Compared to the RW reference, SRRW features a slow decay of *P*_*c*_ at the sub-Mb scale, which is again in consistence with experimental findings^6^, and as revealed in Fig. 2N, the contact frequency is higher inside the tree domains. Interestingly, the decay exponent, −0.75, of intra-tree *P*_*c*_ coincides with the value reported by Hi-C experiments for contacts inside TADs^7^. Comparing intra- and inter-tree contacts, our model predicts that, in single cells, the genomic isolation across tree domains can be as large as 10-fold near the 1Mb range, much higher than the puzzlingly small 2-fold isolation across TADs at the population-average level observed in Hi-C experiments^36,37^. The sufficient isolation at the single-cell level predicted by our model is provided by the elongated inter-tree backbone segments (∼5% of the modeled chromatin), which are expected to be openly accessible and transcriptionally active. This prediction naturally explains the experimental finding that TAD boundaries are active in transcription^38^. Since tree domains can further cluster, and a large tree domain itself has a structural hierarchy, the contact pattern at the TAD (Mb) scale is therefore hierarchical (Fig. S2). Whether this hierarchy extends to large scales is a matter of debate^34,39,40^. While SRRW predicts compartmentalized organizations at different scales, the arrangement of the compartments at the largest scale (>10Mb) (Fig. 2K) is non-hierarchical and can be unraveled to a polymeric string of tree domains (represented by balls) as shown in Fig. 2P. This “secondary” structure of our modeled chromatin highlights the functional importance of the tree domains, which exert strong topological constraints on the genomic organization at the single-cell level.

## Tree domains and larger compartments in a 3D forest

The SRRW model predicts that the topological domains at the single-cell level have random tree structures. Such tree domains are amalgams of chromatin nanoclusters and loops (likely supercoiled) at the kb-to-Mb scale and serve as building blocks of larger compartments. Collectively these tree domains form a “3D forest” within a chromosome territory. A 3D map of tree domains of a typical chromatin structure predicted by SRRW is shown in Fig. 4A, with the color corresponding to the genomic size of each domain. The physical sizes of the tree domains measured by their radius of gyrations (*R*_*g*_) are positively correlated with their genomic sizes but with considerable dispersion (Fig. 4B). The peak *R*_*g*_ of tree domains is around 70nm as shown in Fig. 4C. The genomic size distribution of the tree domains is shown in Fig. 4D, featuring a broad spectrum and an abundance of small domains. The average genomic size is around 50kb, while most of the DNA is packed in tree domains around 300kb, resembling the typical size of TAD at Hi-C level^36^. Interestingly, recent high-resolution TAD analyses revealed a large population of small TADs of tens of kb^41,42^, which aligns well with our tree-domain analysis at the single-cell level. It is worth noting that the genomic size distribution of tree domains is predicted to be strongly non-Gaussian with a heavy tail. As a result, the median size is much smaller than the average size. It is interesting that significant median-average discrepancy has also been reported for TADs whose median size is about 185kb^6^ and average size about 800kb^43^. While the values would depend on the Hi-C resolution, the heavy-tailed domain statistics has been clearly shown in recent sub-kb Hi-C analysis^42^. Our model suggests that such non-Gaussian statistics is a general feature of chromatin, in contrast to normal polymer behavior (Fig. S3).

**Figure 4.**
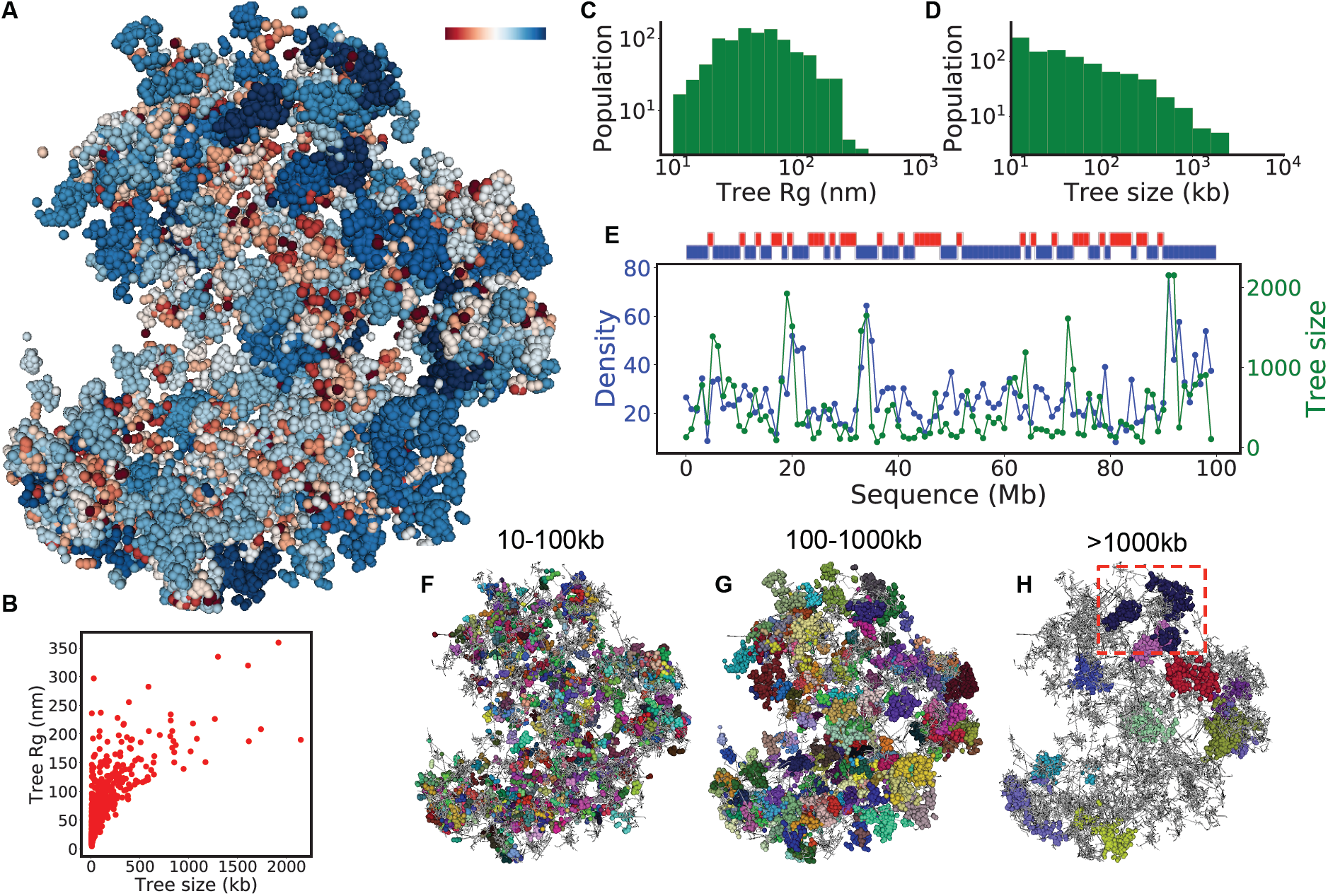
A 3D map of tree domains and their structural statistics. (**A**) Particle representation of the modeled chromatin with tree domains colored according to their genomic sizes (Red: small, Blue: large). (**B**) Scatter plot of the physical size (*R*_*g*_) of tree domains versus their genomic sizes with each domain being one point. (**C**) The physical size (*R*_*g*_) distribution of tree domains. (**D**) The genomic size distribution of tree domains. (**E**) Local DNA density and averaged tree domain size walking along the sequence of the modeled chromatin with each step being 1Mb. Above the panel, the local density is divided into two groups, showing in red (low density) and blue (high density). (**F-H**) Tree domains marked by random colors in three size groups (10-100kb, 100-1000kb, and above 1000kb). The unmarked structures are shown in grey. An example of sub-domains inside a tree domain is highlighted in H.

To reveal the regulatory effect of tree domains on local DNA accessibility, we investigated the relationship between tree domain size and packing density. Remarkably, we have observed a positive correlation between the genomic size of tree domains and the local packing density walking along the modeled chromatin sequence (Fig. 4E), suggestive of a size-dependent domain activity that is in line with recent finding that active domains are small in genomic size^41^. The more condensed packing of larger tree domains is reflected in a decrease of the *R* scaling factor in the sub-Mb regime (Fig. 2L). Using a dichotomy of low (red) and high (blue) packing densities (low density defined as below 80% of mean), we found a quasi-self-similar density pattern (upper Fig. 4E) along the sequence of our modeled chromatin. Resembling the experimentally observed AB-compartment activity alternation^29^, such pattern reflects an emergence of higher-order compartmentalization above tree domains as implied by the plaid-like matrix in Fig. 2K. Such compartments emerge due to the Lévy-flight-like statistics of the secondary structure of SRRW (Fig. 2P), which is non-Gaussian and inhomogeneous at large length scale. The higher-order compartmentalization, however, does not compromise the physical identities of individual tree domains. As shown in Fig. 4F-H, the tree domains (randomly colored) are physical entities with minimal intermixing. Their hierarchical inner structure also lowers the risk of self-entanglement and knotting at sub-Mb level. These structural properties make tree domains good candidates for the functional modules of chromatin at the single-cell level. The similarities between our single-cell predictions and the population-level Hi-C observations suggest that TADs are the statistical consequence of chromatin folding into tree-like topological domains (Fig. S2).

## A quasi-self-similar heterogeneous packing picture of chromatin

One natural yet important structural consequence of SRRW is that the packing density is highly non-uniform across many length scales. To highlight the multi-Mb compartments, we calculated a 3D density map with the nano-scale density fluctuations filtered out. In Fig. 5A, the compartments are demonstrated in a colored particle representation with larger and darker particles corresponding to higher compartmental DNA density. The cross-sections of the filtered 3D density map in all three directions are displayed in density heat maps (upper panels of Fig. 5A). Zooming into a local fraction of the modeled chromatin (without an applied filter) reveals nanoscale domains that are interconnected by physically extended backbone segments (lower left panel of Fig. 5A), and rich in high-density tree nodes as highlighted by golden particles in the second lower panel of Fig. 5A. Our results are consistent with the experimental observation of widespread chromatin nanoclusters or clutches interspersed by nucleosome-depleted regions^44^, and further suggest that large clutches as large tree nodes host higher-order interactions. Our structural prediction of the spatial separation of open and condensed chromatin domains with the latter being larger clumps is also remarkably in line with recent super-resolution observation where active and repressive histone marks have little co-localization and correlate respectively with small low-density and large high-density DNA regions^45^. Our prediction of the ubiquitous alternation between open and condensed chromatin domains suggests that transcriptional activation and repression are highly coupled, like the two sides of a coin. In particular, the model predicts that segments with higher transcriptional propensity tend to serve as stronger isolators between neighboring domains (Fig. S2). The granular packing structure depicted by SRRW registers large surface area and rich porosity, resulting in a similarly heterogeneous encompassing media that could serve as a nuclear channel system^46^. Such a unique space-filling property of SRRW sets it apart from normal disordered polymers, by greatly widening the local DNA density spectrum as shown in Fig. 5B. This broad density spectrum echoes the recent ChromEMT observation^2^, and emphasizes the fundamental role of disordered chromatin packing in transcriptional regulation and nuclear transport.

**Figure 5.**
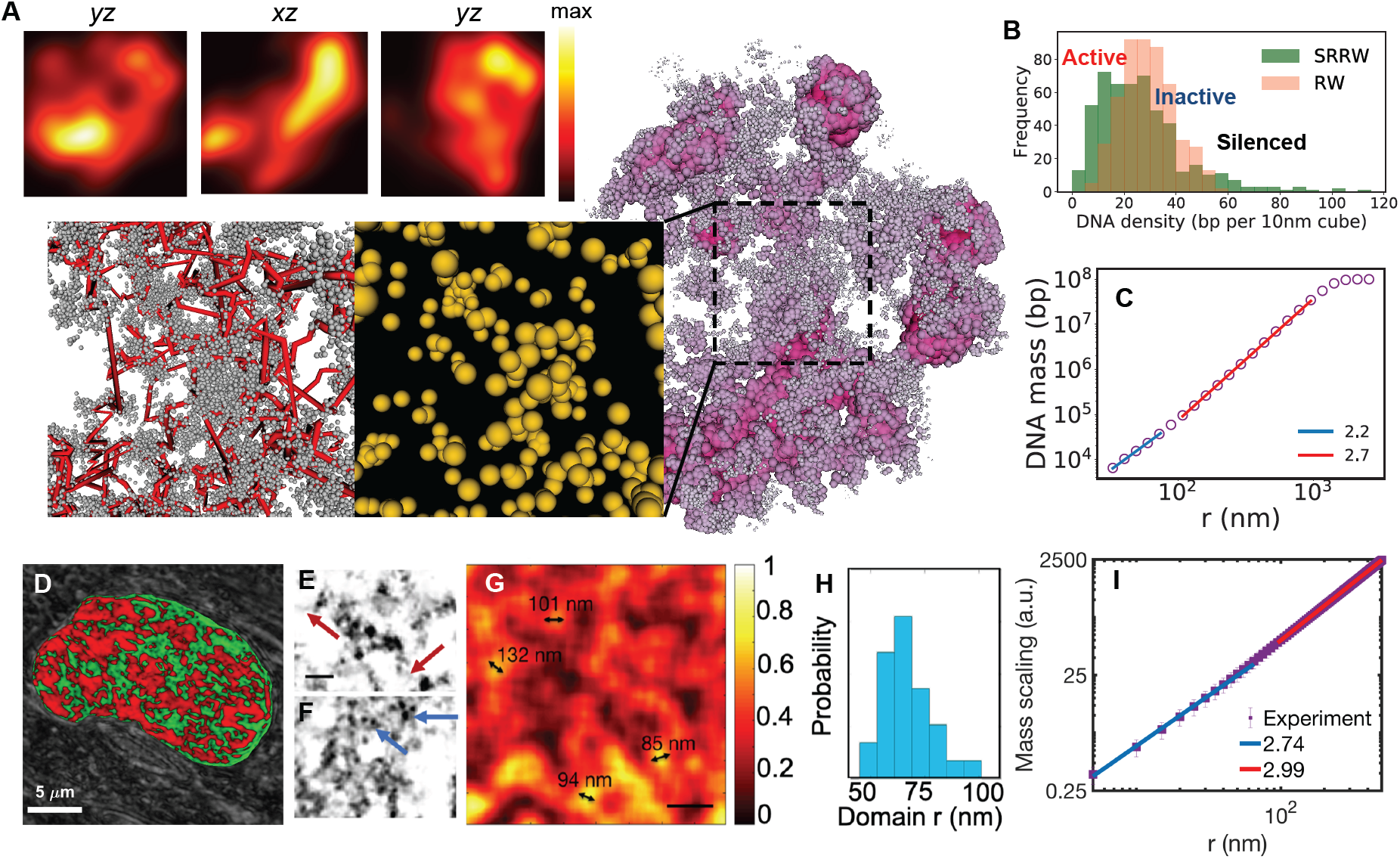
Granular and porous chromatin packing across scales demonstrated by model and experiments. (**A**) Density field representation (with nanoscale fluctuation filtered) of the modeled chromatin and its cross-sections in three orthogonal directions. Zoomed-in panels show the nanoscale packing of the modeled chromatin (without filter). Backbone segments are represented in red lines and nanocluster tree nodes as particles with the volume being proportional to the genomic size. (**B**) Local DNA density spectrum sampled by walking along the modeled chromatin with a probe of 150nm radius (green). A reference is sampled from a RW (red). (**C**) DNA mass scaling of the modeled chromatin (sampled over 1000 SRRW trajectories). (**D**) Typical PWS image of life cell showing compartments of heterogeneous chromatin packing (colored in red). (**E, F**) Linker DNA (red arrows) and individual nucleosomes (blue arrows) captured by ChromSTEM (scale bar: 30nm). (**G**) Chromatin volume concentration based on the convolution of ChromSTEM image (scale bar: 200nm). (**H**) Histogram of chromatin domain size distribution based on the chromatin volume concentration. (**I**) DNA mass scaling based on ChromSTEM.

The existence of two modes of chromatin organization (tree domains and chromatin compartments) leads to an increase in the mass-scaling factor from small to large length scales as shown in Fig. 5C. Such a trend is concordant with the biphasic finding from small angle neutron scattering^24^. Since neutron scattering does not distinguish nuclear molecules, we carried out a ChromSTEM method that focuses on the DNA mass distribution. Combining electron microscopy and DNA staining, ChromSTEM captures nucleosome-level chromatin nanostructure (Fig. 5E, F). Consistent with recent ChromEMT observations, no 30nm fibers were observed. The DNA mass-scaling readout of ChromSTEM (Fig. 5I) confirmed the biphasic behavior predicted by the SRRW model. Despite the slow mass scaling at the small length scale, the tree topology predicted by our model enables high contact frequency within the nanoscale chromatin domains (Fig. 2N), reconciling the contact-structure conundrum present in non-branching polymer models of the higher-order chromatin organization. A DNA density map based on ChromSTEM clearly reveals a granular and porous packing of DNA as shown in Fig. 5G, in line with our prediction. Notably, the peak radius of the chromatin domains revealed by ChromSTEM is around 70 nm (Fig. 5H), coinciding with the peak *R*_*g*_ of tree domains predicted by SRRW (Fig. 4C). The excellent agreement between our model and nanoscopic imaging strongly suggests the existence of functional chromatin modules with minimal intermixing at the sub-Mb level in single cells. From the nanoclusters (clutches) to the tree domains, and to the larger-scale compartments, our model predicts that chromatin has persistent packing heterogeneity across many length scales, hence resembling mass fractals (granular and porous in a quasi-self-similar way) at the single-cell level. To further investigate chromatin packing in live cells, we next performed optical imaging using PWS microscopy^3^. As a label-free technique, PWS microscopy non-invasively measures the intracellular macromolecular arrangement and detects mass-fractal-like structures. As shown in Fig. 5D, PWS microscopy clearly reveals that chromatin is rich in compartments of heterogeneous and mass-fractal-like packing in the live-cell nucleus.

## Emergence of a universal folding principle

As a minimal model, SRRW uses a single folding parameter *α* to describe the collective conformational freedom of chromatin. Here we show that tuning *α* leads to structural alternations across many scales. At the tree domain level, a small *α* promotes the formation of large tree domains and concomitantly more nanoclusters or clutches^44^ as tree nodes, whereas a large *α* reduces the branching and clustering of the modeled chromatin. This is shown in Fig. 6A, where we marked the five largest tree domains in red and the tree nodes larger than 40kb in yellow, for typical architectures at *α*=1.1 and *α*=1.3. At the compartment level, small *α* results in a small population of large compartments while large *α* favors a large collection of small ones. This can be seen in typical physical distance matrices (Fig. 6B, C), where fine plaid-like pattern appears when large packing domains are repressed at large *α*. Recall that from Fig. 2I-K, we have predicted the existence of TAD-like and AB-compartment-like organizations at the single-cell level. Here our structural analysis further suggests a coupling between the repression of packing domains and the finer compartmentalization in single cells. Our prediction finds an interesting analog with the Hi-C observation where weakening TADs unmasks a finer compartment structure at the population-level^40^. Such counteraction between the two modes of chromatin organization can be intuitively understood in the framework of SRRW since the formation of tree domains driven by higher-order interactions imposes hierarchical topological constraints on heterogeneous chromatin segments and suppresses large-scale phase separation.

**Figure 6.**
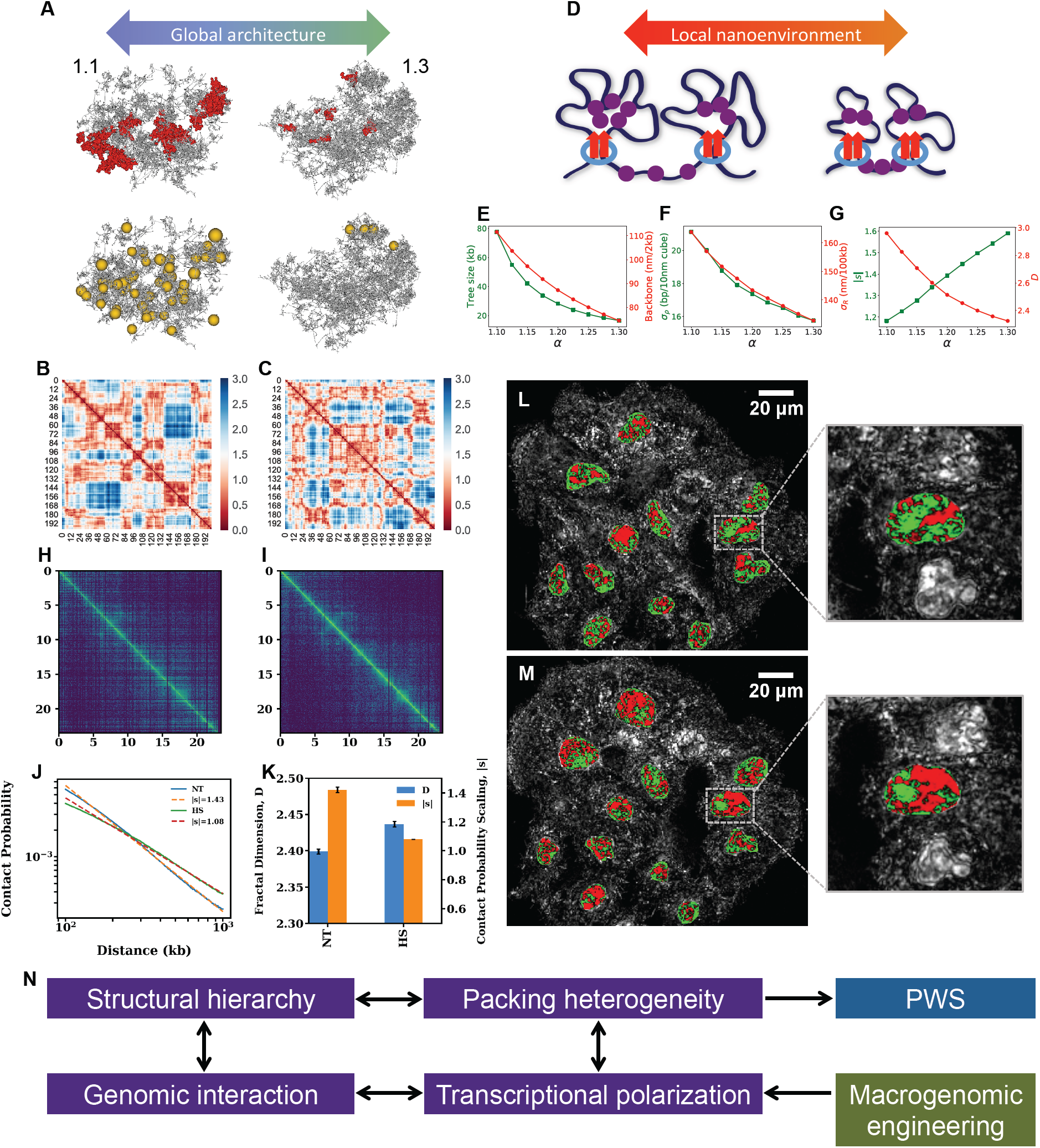
Coupling between chromatin properties during structural alternation. (**A**) Global architectural alternation as *α* changes. Left: *α*=1.1, right: *α*=1.3. For each structure, the five largest tree domains are shown in red in the first row, and the tree nodes (nanoclusters) larger than 40kb in yellow in the second row. (**B, C**) Physical distance maps at *α*=1.1 (left) and *α*=1.3 (right). (**D**) Schematic representation of two different nanoenvironments at small (left) and large (large) *α*. (**E**) Tree domain sizes and backbone openness as functions of *α*. (**F**) How contact scaling and effective mass scaling for 100kb-1Mb modeled chromatin segment depend on *α*. (**G**) Structural heterogeneity of the modeled chromatin measured by the local DNA density variance and mean square distance variance of local DNA segments (sampled over 1000 conformations). (**H, I**) Contact matrices of Kc167 cells exposed to heat shock (HS) and at normal temperature (NT). Entire chromosome scale in the units of Mb. (**J**) Contact probability curves for NT and HS for the 100kb-1Mb range. (**K**) Trends of contact probability scaling change and heterogeneity change after heat shock based on Hi-C analysis and PWS measurements. (**L, M**) PWS live-cell images of HCT116 cells at NT and HS conditions. (**N**) The coupling between different chromatin properties and the idea of macrogenomic engineering for whole-scale transcriptional modulation.

The interplay between the two layers of chromatin organization allows a global architectural change to affect local nanoenvironments where transcription happens. As schematically shown in Fig. 6D, a global architecture rich in large tree domains (small *α*) has more polarized local DNA accessibility and hence transcription activity. While there is more DNA densely packed in large tree domains, the backbone chromatin segments connecting the tree domains are expected to become more open when the domain sizes increase. Numerical analyses of the average tree domain size and backbone openness (physical extent) as functions of *α* have confirmed this picture and are shown in Fig. 6E. Our analysis also shows that polarization of chromatin accessibility is coupled with DNA density heterogeneity (Fig. 6F). To study the local contact-structure relation, we analyzed the contact frequency scaling and the *R* scaling for the modeled chromatin segments of 100kb-1Mb. We fitted the scaling curves to power laws and extracted scaling factors (Fig. S4). An effective mass-scaling factor *D* defined as the inverse of *R* scaling factor measures the mass scaling based on consecutive DNA mass with definite segment length. Fig. 5G shows that *D* decreases as *α* goes up, anti-correlated with the contact scaling *s*, but correlates with the local packing heterogeneity. The coupling between mass scaling and structural heterogeneity suggests, again, that chromatin compartments can be approximated as a mass fractal with its structural heterogeneity quantified by a fractal dimension *D*. It merits a note that our prediction of higher chromatin heterogeneity fostering more genomic contacts contrasts the behavior of a fractal chain (Fig. 1A). The reason why chromatin resembles 3D mass fractal rather than 1D fractal chain is due to the prevailing clusters and branches that lead to a heterogeneous and porous structure.

The above theoretical analysis suggests that during chromatin structural alternation, which could happen when cell responses to stress, the perturbations of many chromatin properties are coupled. Here we test this prediction using heat shock as model experimental system. We analyze the contact maps (Fig. 6H, I) and contact probability curves (Fig. 6J) at normal temperature and exposed to heat shock using the publicly available Hi-C data on Kc167 *Drosophila* cells (GEO access code GSE63518)^47^. In line with literature^47^, we noticed a decrease in contact probability scaling with the heat shock compared to the control condition, which signifies an increase in long-range contacts in Kc167 cells subjected to heat shock. We have carried out live-cell PWS measurements on HCT116 cells to study the effect of heat shock on the packing heterogeneity of chromatin. As shown in Fig. 6L, M, we observed that temperature stress led to a higher PWS signal that means more heterogeneous packing of live-cell chromatin. Together, we found that the contact probability scaling factor *s* and the average mass-fractal-dimension *D* are anti-correlated as shown in Fig. 6K, which is consistent with the prediction of our model (Fig. 6G). Moreover, our model suggests that formation of long-range genomic contacts is coupled with promotion of high-density chromatin clusters, in line with the Hi-C report that chromatin forms more polycomb-silenced heterochromatin while increasing its self-interaction frequencies during heat shock^47^. Altogether, our theoretical and experimental results suggest that structural hierarchy, packing heterogeneity, genomic interaction, and transcriptional polarization are intimately coupled during epigenetic reconfiguration (Fig. 6N), which supports the idea of using macrogenomic engineering^4^ to treat system diseases such as cancer^48^.

## Discussion and conclusion

In this paper, we asked the question of how chromatin folds to achieve simultaneously high self-interacting frequency and high space-filling heterogeneity. While there is no lack of 3D models^49,25,8,17,50^ with conflicting single-cell level predictions developed to explain the population-level Hi-C contact maps, the packing heterogeneity has received less attention and remained poorly understood. Nevertheless, a few lines of recent experimental evidences^2,4,20,46^ have converged to suggest that chromatin packing heterogeneity determines the functional accessibility and activity of interphase DNA. Here we integrated modeling and imaging to understand 3D genome at the single-cell level. We have carried out ChromSTEM and PWS measurements to quantify the heterogeneity of chromatin packing across different length scales. At the nano-scale, concurring with recent study^2^, we did not observe the 30nm fiber, and instead revealed a granular packing structure with prevailing nano-domains of an peak radius around 70nm (Fig. 5H). To characterize chromatin packing at larger scales and in live cells, we have conducted PWS microscopy to reveal that heterogeneous packing compartments exist throughout the nucleus (Fig. 5D). Our multi-scale imaging suggests a quasi-self-similar picture of chromatin packing. Using heat shock as an example, we demonstrated that the average heterogeneity characterized by a mass-fractal dimension is sensitive to environmental change and could serve as an indicator of chromatin folding state (Fig. 6L, M).

There is a subtle yet important topological difference between a fractal chain and a mass fractal, as the former does not branch. We theoretically proved^23^ that for 1D chain, there is a trilemma between high packing heterogeneity, high contact frequency, and self-similarity. This means that the presumed non-branching topology and the strict scale-invariance of chromatin must be broken to unify high packing heterogeneity and contact frequency, hence necessitating distinct folding modes at different scales. In this light, we carried out ChromSTEM to find a transition in the mass scaling near 100nm (Fig. 5I), close to the peak domain radius. Note that the scale-dependent power laws have been also reported in literature for chromatin contact frequency^7^ and mean square distance^51^. To understand the scale-dependent packing modes, we introduced a self-returning random walk (SRRW) model, which generates tree-like topological domains with hierarchical structures (Fig. 1). The tree domains pack local genomic DNA into nested loops and nanoclusters with high contact frequency at the sub-Mb scale, and constitute mass-fractal-like compartments at the multi-Mb scale (Figs. 2, 4). We predict that an open chromatin backbone connects and isolates the tree domains, rendering highly porous chromatin structure, flexible higher-order folding, and irregularly shaped chromosome territory (Fig. 5A-C). Our model agrees well with our experiments’ demonstrating (Fig. 5D-I) a heterogeneous chromatin structure across many length scales. Moreover, the model predicts an anti-correlation between contract frequency scaling and packing heterogeneity, consistent with our experimental observation of heat shock response (Fig. 6).

It is interesting that our model predicts TAD-like and AB-compartment-like patterns on a distance matrix at the single-cell level (Fig. 2I-K), resembling the Hi-C contact patterns^6^ at the population level. This resemblance becomes more intriguing, since the model further predicts a counteraction between the two single-cell organizational modes (Fig. 6B, C), reminiscent of the similar phenomenon at the population level reported in Hi-C experiments^40^. This suggests that well-organized 3D structures such as domains and compartments are intrinsic chromatin characteristics at the single-cell level rather than merely statistical patterns of large population (Fig. S2). Furthermore, our model predicts that the 3D genome is more structured than that suggested by population-level contact patterns. Beyond the average 2-fold isolation associated with TADs^36,37^, tree domains allow single-cell chromatin structure to realize 5-to 10-fold probability difference between the intra- and inter-domain contacts at the 100kb-1Mb level (Fig. 2N). Compared to the diffuse and intermixing loops predicted by the loop extrusion hypothesis^8^, our model depicts more compacted and isolated physical entities in single cells (Fig. 3), rich in spontaneous higher-order interactions as revealed by recent super-resolution experiment^9^. In line with recent Hi-C reports^41,42^, statistical analysis of our modeled chromatin suggests that there are prevailing small domains at the single-cell level, followed by a heavy tail of larger ones (Fig. 2D, F, F, G). Our model further predicts a correlation between local DNA density and domain size (Fig. 2E), which lends explanation to the experimental finding that small domains are on average more active than large ones^41^. Lastly, our prediction that tree domains are isolated by easily accessible chromatin backbone segments (Fig. 2P, Fig. 3) is consistent with the observation that TAD boundaries are enriched in active genes^38^.

It is remarkable that by resolving the contact-structure paradox (Fig. 1), a minimal model is able to pivot a wide array of chromatin features including untangled compaction^21^, disordered morphology^2^, non-globular territory^29^, porous nuclear medium^46^, biphasic chromatin mass scaling^24^, abundance of nanoclusters or clutches^44^, non-Gaussian domain size distribution^41,42^, size-dependent domain activity^41^, active domain boundaries^38^, higher-order interactions^9^, and multiple layers of chromatin organization^40^. This suggests an emerging picture that, on top of the first-order genomic structure, namely the linear DNA double helix for data reproduction, our genome has evolved to physically adopt virtual tree data structure (Fig. 1F) for its higher-order functional modules to better organize the enormous genomic information. In this regard, functional optimization may be a better perspective to understand chromatin folding than polymer physics. In particular, we argue that forming a string of tree-like topological domains connected by open backbone segments has functional advantages in (1) proper organization of genomic contacts, (2) packing-based regulation of transcription, (3) transport and accommodation of nuclear proteins, and (4) transition between interphase and mitosis.

Proper genomic contacts based on chromatin folding are critical to cell function. In stark contrast to the equilibrium globule reference (Fig. 2F-H), we have shown that our model promotes short-range contacts and suppresses long-range (multi-Mb and above) contacts. Since longer-range contacts have more combinatorial possibilities, this suggests that chromatin folding can significantly lower the information entropy of genomic interaction, despite the observed “disordered” polymer structure^2^. Such ordered genomic organization not only enhances the local regulatory contacts within hierarchical domains but also restricts random long-range contacts that could lead to aberrant functions (note that errant long-range genomic interactions are hallmarks of cancer^52–54^).

The identity of cell depends on the coordination of highly active and strictly silenced genes. A widened spectrum of local DNA density along the sequence as predicted by our model (Fig. 5B) allows for a polarized accessibility of genome to ensure that particular genes are active or silenced. On the other hand, the quasi-self-similar packing picture revealed by our experiments and model (Fig. 5) is advantageous in supporting fast transport of nuclear proteins of varying sizes. Such porosity across many length scales can also accommodate the experimentally observed clusters and condensates from tens of nanometer to micron scale that are comprised of different transcription factors, DNA-processing enzymes, and chromatin architectural proteins^36,55,56^.

It is worth noting that in our model, the majority of chromatin is packed into tree domains whereas only about 5% of interphase DNA is exposed in the backbone segments that are physically extended and of high transcriptional competence. It is this minority of the heterogeneous biopolymer that creates most of the low DNA density regions in the porous chromatin structure. Recent in vitro experiment^57^ confirmed a condensin loop extrusion mechanism^58^ on naked DNA, which is strictly one-sided and has been argued^59^ to be far from sufficient for mitotic condensation. Nevertheless, based our prediction that only about 5% of the DNA swells the porous chromosome territory, the one-sided condensin extrusion on this extended backbone of chromatin could have a considerable effect of collapsing interphase DNA into mitotic form. More importantly, such backbone extrusion could preserve the folding memory of chromatin into the topological information of tree domains even though most of the Hi-C detectable^60^ spatial proximity information between loci is inevitably lost during the transition from interphase to mitosis. In this way, our folding picture allows reconciliation between mitotic condensation and folding memory inheritance. Consistently, recent super-resolution single nucleosome tracking suggested the existence of chromatin domains throughout the cell cycle^61^. Remarkably, the reported peak radius of the domains is about 80nm, similar to our ChromSTEM measurement and model’s prediction.

The formation of tree domains predicted by our model allows the functional modules of chromatin to have topological identities (a robust way to store information). However, such topological domains cannot be self-assembled without energy input (note SRRW as a folding algorithm is not a memory-less Markov process and cannot happen at thermodynamic equilibrium). Nevertheless, a tree-like topology of chromatin segment is not totally unexpected since looping and supercoiling can effectively branch the biopolymer. Given the structural complexity and heterogeneity of tree domains predicted by SRRW, we expect the folding process to involve a diversity of molecular mechanisms to act in concert under the guidance of genetic and epigenetic codes. These could include transcription-induced supercoiling^11–14^, phase separation^15–17^, long non-coding RNA^62^, DNA-mediated charge transfer^63^ and the putative cohesive-CTCF loop extrusion^8^. In particular, DNA-mediated charge transfer could enable fast long-range communication between responsible architectural proteins in a transcription-sensitive way, whereas long non-coding RNA could allow slow but more specific loci communication and recruitment of folding agents. Given the hierarchical nature of the tree topology, we expect both the assembly and disassembly of the tree domains to be hierarchical processes, meaning that small tree domains can group into large ones where they become sub-domains and vice versa like chemical reactions. During such grouping and ungrouping processes, backbone segments can be absorbed into new tree domains and new backbone segments can be exposed. For a steady state when the cell is not preparing for division, we expect the overall statistics (Fig. 4), i.e., the tree size distribution, the degree of structural hierarchy, and the portion of backbone segments to be stable. Like the inheritance of non-coding DNA (>90% of genome), maintaining physically extended backbone segments (∼5% of genome) and their low DNA density local environment inside the highly crowded nucleus may seem uneconomical, since energy needs to be consumed against the conformational entropy of chromatin that tends to homogenize the DNA concentration. This puzzle could be explained by recent experimental finding that droplet of nuclear proteins initialized by DNA binding mechanically dispels heterochromatin and promotes the formation of low DNA density region^16^. This picture is consistent with another recent observation that the total euchromatin density including proteins is only 1.5-fold lower than that of the surrounding heterochromatin, despite the 5.5-to 7.5-fold DNA density difference^64^. Altogether, the formation of both tree domains and chromatin backbone in vivo is not only functionally desirable but also mechanistically possible.

The fact that many major chromatin features can be explained by an abstract folding algorithm indicates the existence of universal principles for chromatin to functionally fold and efficiently explore the genomic landscape. Our results suggest a global coupling between different chromatin properties including domain hierarchy, nanocluster size distribution, backbone openness, packing heterogeneity, genomic interaction, and transcriptional polarization, which means a significant dimensionality reduction during chromatin folding due to the emergence of collective folding parameters or susceptibilities (*α* in SRRW for example). Such folding picture sheds new lights into the physics of epigenetics^65–67^ and chromatin plasticity^68^, and stresses the importance of understanding 3D genome from a data structure point of view (Fig. 1F) on top of polymer physics, since chromatin folding is optimizing biological functions rather than minimizing thermodynamic free energy. The unexpected positive correlation between genomic contact scaling and DNA packing heterogeneity of living chromatin under heat shock is one example.

In summary, we have combined theoretical and experimental efforts to understand chromatin topology, statistics, scaling and their couplings at the single-cell level. Our multi-scale results, from kb-level nanoclusters to 100Mb-level chromosome territories, provided an integrative view of higher-order chromatin folding as an alternative thinking to the classical 30nm-fiber-based picture. Our prediction that chromatin folds into tree-like topological domains connected by an active backbone explains and reconciles a wealth of exotic properties of this living biopolymer that are alien to the common sense of polymer physics. It is remarkable that despite the distinct folding modes at different scales, chromatin is able to maintain alternation between active and inactive states across many scales both in space and on DNA sequence, in a quasi-self-similar manner. The non-Gaussian folding statistics and global coupling between chromatin properties suggest that interphase DNA explores the great genomic landscape as a complex network rather than a simple polymer. The possibility of chromatin having tree data structures and universal folding principles opens an exciting new paradigm to understand genomic organization and presents many new questions the answering of which would require collaborations between experimentalists and theorists from different fields. We hope our insights in this paper could inspire future interdisciplinary efforts on this grand challenge of life science.

## Method summary

### Model

The folding algorithm of SRRW is given in the main text. Here we summarize some parameters we used in the model and data analysis. The smallest step size in the model is 30nm. For local DNA density analysis we use a probe of 150nm radius. For contact frequency calculation we choose a contact criterion to be two loci being within 45nm. The single-cell level modeling and analyses are done at *α*=1.15. The local and global cutoffs in this case are 24.85 and 57.15 in reduced units. Population-level modeling and analyses are done over 1000 independent samples.

### ChromEM staining with STEM tomography and TEM imaging (ChromSTEM)

ChromEM staining was employed to label the DNA in the A549 cell and BJ cell nucleus as previously described^2^. The cells were fixed in 2.5% EM grade glutaraldehyde and 2% paraformaldehyde (EMS, USA) in 1x sodium cacodylate buffer for 20 min at room temperature and 1h on ice. All the following steps were conducted on ice or a customized cold-stage with temperature control, and the reagents used in the protocol were pre-chilled in the fridge or on ice. After fixation, the cells were stained with DRAQ5^TM^ (Theromo Fisher, USA) that intercalates ds-DNA and bathed in 3,3’-diaminobenzidine (DAB) solution (Sigma Aldrich, USA). An inverted microscopy (Eclipse, Nikon, Japan) was used for photo-bleaching. Each spot was photo-bleached for 7 min by epi illumination with a 100x objective. Osmium ferrocyanide was used to further enhance contrast of the DNA and standard Durcupan^TM^ resin (EMS, USA) embedding was carried out. After curing for 48 hours at 60 °C, ultra-thin sections were prepared by an ultramicrotome (UC7, Leica, USA) and deposited onto plasma-treated TEM grid with formar-carbon film (EMS, USA). To investigate the CVC and radius of the chromatin nano-domains, dual-axis STEM HAADF tomography was performed on the 100 nm ultra-thin section of A549 cell nucleus with 10 nm colloidal gold fiducial markers. The sample was tilted from −60° to 60° with 2° step along the two perpendicular axes respectively. IMOD^69^ was employed to align the tilt series, and the tomography reconstruction was conducted in TomoPy^70^ using a penalized maximum likelihood algorithm (PLM-hybrid) for each axis. The combination of tomograms was performed IMOD to further suppress the influence of missing cone. Furthermore, to estimate the average cluster size for the whole nucleus across multiple cells experimentally, mass-scaling analysis of TEM projection images on ultra-thin sections of BJ cells was performed. The TEM contrast was converted to mass-thickness by the Beer’s law prior to the mass-scaling analysis.

### Live-cell Partial Wave Spectroscopic (PWS) Microscopy

Live-cell Partial Wave Spectroscopic microscopy measurements were performed as previously^3^. Briefly, cells were seeded onto size-0 glass-bottom 35mm dishes in McCoy’s 5A medium (Lonza) media supplemented with 10% FBS and 1X Penicillin-Streptomycin at least 48 hours prior to imaging at a concentration of 50,000 cells per ml and cultured at 37 °C with 21% O2 and 5% CO2. Cells were then imaged using a broad spectrum LED with illumination focused onto the sample plane using a 63x or 100x objective as indicated. Backscattered light was collected with spectral filtration performed utilizing a liquid-crystal tunable filter (LCTF, CRi, Woburn, MA; spectral resolution 7nm) collecting sequential monochromatic images between 500nm and 700nm. These monochromatic images were projected onto a CMOS camera (ORCA-Flash 4.0 v2, Hamamatsu City, Japan) producing a three-dimensional image cube. Variations in the backscattered intensity were analyzed at each pixel. Mean spectral variations for manually segmented nuclei were analyzed for the following conditions-cells incubated at 37 °C (n=1046), and cells incubated at 42 °C for 1 hour (n=1080). With the exception of heat-shock experiments, all measurements were performed at 37 °C with 21% O2 and 5% CO2 using a stage-top environmental chamber.

## Acknowledgements

We gratefully acknowledge funding from National Science Foundation: Biol & Envir Inter of Nano Mat 1833214, EFRI research projects 1830961, and from the National Institutes of Health: National Cancer Institute R01 CA228272 and R01 CA225002. We thank members of the Szleifer and Backman laboratories for their insights and comments.

## Author contributions

K.H., V.B., and I.S. conceived the project. K.H. developed the theory and model. Y.L. performed the ChromSTEM experiments and analyzed the data. V.A. and A.E. performed PWS experiments and analyzed the data. R.V. analyzed the publicly available Hi-C data on heat shock. K.H. wrote the original draft and all authors reviewed and edited the manuscript. V.B. and I.S. secured the funding and supervised the project. The authors have no competing interests.

## References

(1) Finch, J.; Klug, A. Solenoidal Model for Superstructure in Chromatin. Proc Natl Acad Sci USA 1976, 73 (6), 1997–1901.

(2) Ou, H. D.; Phan, S.; Deerinck, T. J.; Thor, A.; Ellisman, M. H.; O’Shea, C. C. ChromEMT: Visualizing 3D Chromatin Structure and Compaction in Interphase and Mitotic Cells. Science 2017, 357 (6349), eaag0025. https://doi.org/10.1126/science.aag0025.

(3) Almassalha, L. M.; Bauer, G. M.; Chandler, J. E.; Gladstein, S.; Cherkezyan, L.; Stypula-Cyrus, Y.; Weinberg, S.; Zhang, D.; Thusgaard Ruhoff, P.; Roy, H. K.; et al. Label-Free Imaging of the Native, Living Cellular Nanoarchitecture Using Partial-Wave Spectroscopic Microscopy. Proc. Natl. Acad. Sci. 2016, 113 (42), E6372–E6381. https://doi.org/10.1073/pnas.1608198113.

(4) Almassalha, L. M.; Bauer, G. M.; Wu, W.; Cherkezyan, L.; Zhang, D.; Kendra, A.; Gladstein, S.; Chandler, J. E.; VanDerway, D.; Seagle, B.-L. L.; et al. Macrogenomic Engineering via Modulation of the Scaling of Chromatin Packing Density. Nat. Biomed. Eng. 2017, 1 (11), 902–913. https://doi.org/10.1038/s41551-017-0153-2.

(5) Lieberman-Aiden, E.; van Berkum, N. L.; Williams, L.; Imakaev, M.; Ragoczy, T.; Telling, A.; Amit, I.; Lajoie, B. R.; Sabo, P. J.; Dorschner, M. O.; et al. Comprehensive Mapping of Long-Range Interactions Reveals Folding Principles of the Human Genome. Science 2009, 326 (5950), 289–293. https://doi.org/10.1126/science.1181369.

(6) Rao, S. S. P.; Huntley, M. H.; Durand, N. C.; Stamenova, E. K.; Bochkov, I. D.; Robinson, J. T.; Sanborn, A. L.; Machol, I.; Omer, A. D.; Lander, E. S.; et al. A 3D Map of the Human Genome at Kilobase Resolution Reveals Principles of Chromatin Looping. Cell 2014, 159 (7), 1665–1680. https://doi.org/10.1016/j.cell.2014.11.021.

(7) Sanborn, A. L.; Rao, S. S. P.; Huang, S.-C.; Durand, N. C.; Huntley, M. H.; Jewett, A. I.; Bochkov, I. D.; Chinnappan, D.; Cutkosky, A.; Li, J.; et al. Chromatin Extrusion Explains Key Features of Loop and Domain Formation in Wild-Type and Engineered Genomes. Proc. Natl. Acad. Sci. 2015, 112 (47), E6456–E6465. https://doi.org/10.1073/pnas.1518552112.

(8) Fudenberg, G.; Imakaev, M.; Lu, C.; Goloborodko, A.; Abdennur, N.; Mirny, L. A. Formation of Chromosomal Domains by Loop Extrusion. Cell Rep. 2016, 15 (9), 2038–2049. https://doi.org/10.1016/j.celrep.2016.04.085.

(9) Bintu, B.; Mateo, L. J.; Su, J.-H.; Sinnott-Armstrong, N. A.; Parker, M.; Kinrot, S.; Yamaya, K.; Boettiger, A. N.; Zhuang, X. Super-Resolution Chromatin Tracing Reveals Domains and Cooperative Interactions in Single Cells. Science 2018, 362 (6413), eaau1783. https://doi.org/10.1126/science.aau1783.

(10) Szabo, Q.; Jost, D.; Chang, J.-M.; Cattoni, D. I.; Papadopoulos, G. L.; Bonev, B.; Sexton, T.; Gurgo, J.; Jacquier, C.; Nollmann, M.; et al. TADs Are 3D Structural Units of Higher-Order Chromosome Organization in *Drosophila*. Sci. Adv. 2018, 4 (2), eaar8082. https://doi.org/10.1126/sciadv.aar8082.

(11) Kouzine, F.; Gupta, A.; Baranello, L.; Wojtowicz, D.; Ben-Aissa, K.; Liu, J.; Przytycka, T. M.; Levens, D. Transcription-Dependent Dynamic Supercoiling Is a Short-Range Genomic Force. Nat. Struct. Mol. Biol. 2013, 20 (3), 396–403. https://doi.org/10.1038/nsmb.2517.

(12) Naughton, C.; Avlonitis, N.; Corless, S.; Prendergast, J. G.; Mati, I. K.; Eijk, P. P.; Cockroft, S. L.; Bradley, M.; Ylstra, B.; Gilbert, N. Transcription Forms and Remodels Supercoiling Domains Unfolding Large-Scale Chromatin Structures. Nat. Struct. Mol. Biol. 2013, 20 (3), 387–395. https://doi.org/10.1038/nsmb.2509.

(13) Corless, S.; Gilbert, N. Effects of DNA Supercoiling on Chromatin Architecture. Biophys. Rev. 2016, 8 (3), 245–258. https://doi.org/10.1007/s12551-016-0210-1.

(14) Kim, S. H.; Ganji, M.; Kim, E.; van der Torre, J.; Abbondanzieri, E.; Dekker, C. tDNA Sequence Encodes the Position of DNA Supercoils. eLife 2018, 7. https://doi.org/10.7554/elife.36557.

(15) Sabari, B. R.; Dall’Agnese, A.; Boija, A.; Klein, I. A.; Coffey, E. L.; Shrinivas, K.; Abraham, B. J.; Hannett, N. M.; Zamudio, A. V.; Manteiga, J. C.; et al. Coactivator Condensation at Super-Enhancers Links Phase Separation and Gene Control. Science 2018, 361 (6400), eaar3958. https://doi.org/10.1126/science.aar3958.

(16) Shin, Y.; Chang, Y.-C.; Lee, D. S. W.; Berry, J.; Sanders, D. W.; Ronceray, P.; Wingreen, N. S.; Haataja, M.; Brangwynne, C. P. Liquid Nuclear Condensates Mechanically Sense and Restructure the Genome. Cell 2018, 175 (6), 1481- 1491.e13. https://doi.org/10.1016/j.cell.2018.10.057.

(17) Di Pierro, M.; Cheng, R. R.; Lieberman Aiden, E.; Wolynes, P. G.; Onuchic, J. N. De Novo Prediction of Human Chromosome Structures: Epigenetic Marking Patterns Encode Genome Architecture. Proc. Natl. Acad. Sci. 2017, 114 (46), 12126–12131. https://doi.org/10.1073/pnas.1714980114.

(18) Bianco, S.; Chiariello, A. M.; Annunziatella, C.; Esposito, A.; Nicodemi, M. Predicting Chromatin Architecture from Models of Polymer Physics. Chromosome Res. 2017, 25 (1), 25–34. https://doi.org/10.1007/s10577-016-9545-5.

(19) Kim, J. S.; Backman, V.; Szleifer, I. Crowding-Induced Structural Alterations of Random-Loop Chromosome Model. Phys Rev Lett 2011, 106, 168102. https://doi.org/10.1103/PhysRevLett.106.168102.

(20) Schwartz, U.; Németh, A.; Diermeier, S.; Exler, J. H.; Hansch, S.; Maldonado, R.; Heizinger, L.; Merkl, R.; Längst, G. Characterizing the Nuclease Accessibility of DNA in Human Cells to Map Higher Order Structures of Chromatin. Nucleic Acids Res. 2018. https://doi.org/10.1093/nar/gky1203.

(21) Grosberg, A.; Rabin, Y.; Havlin, S.; Neer, A. Crumpled Globule Model of the Three-Dimensional Structure of DNA. Europhys. Lett. EPL 1993, 23 (5), 373–378. https://doi.org/10.1209/0295-5075/23/5/012.

(22) Mirny, L. A. The Fractal Globule as a Model of Chromatin Architecture in the Cell. Chromosome Res. 2011, 19 (1), 37–51. https://doi.org/10.1007/s10577-010-9177-0.

(23) Huang, K.; Backman, V.; Szleifer, I. Interphase Chromatin as a Self-Returning Random Walk: Can DNA Fold into Liquid Trees? 2018. https://doi.org/10.1101/413872.

(24) Lebedev, D. V.; Filatov, M. V.; Kuklin, A. I.; Islamov, A. K.; Kentzinger, E.; Pantina, R.; Toperverg, B. P.; Isaev-Ivanov, V. V. Fractal Nature of Chromatin Organization in Interphase Chicken Erythrocyte Nuclei: DNA Structure Exhibits Biphasic Fractal Properties. FEBS Lett. 2005, 579 (6), 1465–1468. https://doi.org/10.1016/j.febslet.2005.01.052.

(25) Giorgetti, L.; Galupa, R.; Nora, E. P.; Piolot, T.; Lam, F.; Dekker, J.; Tiana, G.; Heard, E. Predictive Polymer Modeling Reveals Coupled Fluctuations in Chromosome Conformation and Transcription. Cell 2014, 157 (4), 950–963. https://doi.org/10.1016/j.cell.2014.03.025.

(26) Li, G.; Levitus, M.; Bustamante, C.; Widom, J. Rapid Spontaneous Accessibility of Nucleosomal DNA. Nat. Struct. Mol. Biol. 2005, 12 (1), 46–53. https://doi.org/10.1038/nsmb869.

(27) Segal, E.; Fondufe-Mittendorf, Y.; Chen, L.; Thåström, A.; Field, Y.; Moore, I. K.; Wang, J.-P. Z.; Widom, J. A Genomic Code for Nucleosome Positioning. Nature 2006, 442 (7104), 772–778. https://doi.org/10.1038/nature04979.

(28) Chereji, R. V.; Morozov, A. V. Functional Roles of Nucleosome Stability and Dynamics. Brief. Funct. Genomics 2015, 14 (1), 50–60. https://doi.org/10.1093/bfgp/elu038.

(29) Stevens, T. J.; Lando, D.; Basu, S.; Atkinson, L. P.; Cao, Y.; Lee, S. F.; Leeb, M.; Wohlfahrt, K. J.; Boucher, W.; O’Shaughnessy-Kirwan, A.; et al. 3D Structures of Individual Mammalian Genomes Studied by Single-Cell Hi-C. Nature 2017, 544 (7648), 59–64. https://doi.org/10.1038/nature21429.

(30) Cremer, T.; Cremer, M.; Dietzel, S.; Müller, S.; Solovei, I.; Fakan, S. Chromosome Territories – a Functional Nuclear Landscape. Curr. Opin. Cell Biol. 2006, 18 (3), 307–316. https://doi.org/10.1016/j.ceb.2006.04.007.

(31) Sutherland, H.; Bickmore, W. A. Transcription Factories: Gene Expression in Unions? Nat. Rev. Genet. 2009, 10 (7), 457–466. https://doi.org/10.1038/nrg2592.

(32) Bancaud, A.; Huet, S.; Daigle, N.; Mozziconacci, J.; Beaudouin, J.; Ellenberg, J. Molecular Crowding Affects Diffusion and Binding of Nuclear Proteins in Heterochromatin and Reveals the Fractal Organization of Chromatin. EMBO J. 2009, 28 (24), 3785–3798. https://doi.org/10.1038/emboj.2009.340.

(33) Nagano, T.; Lubling, Y.; Stevens, T. J.; Schoenfelder, S.; Yaffe, E.; Dean, W.; Laue, E. D.; Tanay, A.; Fraser, P. Single-Cell Hi-C Reveals Cell-to-Cell Variability in Chromosome Structure. Nature 2013, 502 (7469), 59–64. https://doi.org/10.1038/nature12593.

(34) Tan, L.; Xing, D.; Chang, C.-H.; Li, H.; Xie, X. S. Three-Dimensional Genome Structures of Single Diploid Human Cells. Science 2018, 361 (6405), 924–928. https://doi.org/10.1126/science.aat5641.

(35) Wang, S.; Su, J.-H.; Beliveau, B. J.; Bintu, B.; Moffitt, J. R.; Wu, C.; Zhuang, X. Spatial Organization of Chromatin Domains and Compartments in Single Chromosomes. Science 2016, 353 (6299), 598–602. https://doi.org/10.1126/science.aaf8084.

(36) Hansen, A. S.; Cattoglio, C.; Darzacq, X.; Tjian, R. Recent Evidence That TADs and Chromatin Loops Are Dynamic Structures. Nucleus 2018, 9 (1), 20–32. https://doi.org/10.1080/19491034.2017.1389365.

(37) Dekker, J.; Mirny, L. The 3D Genome as Moderator of Chromosomal Communication. Cell 2016, 164 (6), 1110–1121. https://doi.org/10.1016/j.cell.2016.02.007.

(38) Ulianov, S. V.; Khrameeva, E. E.; Gavrilov, A. A.; Flyamer, I. M.; Kos, P.; Mikhaleva, E. A.; Penin, A. A.; Logacheva, M. D.; Imakaev, M. V.; Chertovich, A.; et al. Active Chromatin and Transcription Play a Key Role in Chromosome Partitioning into Topologically Associating Domains. Genome Res. 2016, 26 (1), 70–84. https://doi.org/10.1101/gr.196006.115.

(39) Fraser, J.; Ferrai, C.; Chiariello, A. M.; Schueler, M.; Rito, T.; Laudanno, G.; Barbieri, M.; Moore, B. L.; Kraemer, D. C.; Aitken, S.; et al. Hierarchical Folding and Reorganization of Chromosomes Are Linked to Transcriptional Changes in Cellular Differentiation. Mol. Syst. Biol. 2015, 11 (12), 852–852. https://doi.org/10.15252/msb.20156492.

(40) Schwarzer, W.; Abdennur, N.; Goloborodko, A.; Pekowska, A.; Fudenberg, G.; Loe-Mie, Y.; Fonseca, N. A.; Huber, W.; Haering, C.; Mirny, L.; et al. Two Independent Modes of Chromatin Organization Revealed by Cohesin Removal. Nature 2017. https://doi.org/10.1038/nature24281.

(41) Ramírez, F.; Bhardwaj, V.; Arrigoni, L.; Lam, K. C.; Grüning, B. A.; Villaveces, J.; Habermann, B.; Akhtar, A.; Manke, T. High-Resolution TADs Reveal DNA Sequences Underlying Genome Organization in Flies. Nat. Commun. 2018, 9 (1). https://doi.org/10.1038/s41467-017-02525-w.

(42) Wang, Q.; Sun, Q.; Czajkowsky, D. M.; Shao, Z. Sub-Kb Hi-C in D. Melanogaster Reveals Conserved Characteristics of TADs between Insect and Mammalian Cells. Nat. Commun. 2018, 9 (1). https://doi.org/10.1038/s41467-017-02526-9.

(43) Dekker, J.; Heard, E. Structural and Functional Diversity of Topologically Associating Domains. FEBS Lett. 2015, 589 (20PartA), 2877–2884. https://doi.org/10.1016/j.febslet.2015.08.044.

(44) Ricci, M. A.; Manzo, C.; García-Parajo, M. F.; Lakadamyali, M.; Cosma, M. P. Chromatin Fibers Are Formed by Heterogeneous Groups of Nucleosomes In Vivo. Cell 2015, 160 (6), 1145–1158. https://doi.org/10.1016/j.cell.2015.01.054.

(45) Xu, J.; Ma, H.; Jin, J.; Uttam, S.; Fu, R.; Huang, Y.; Liu, Y. Super-Resolution Imaging of Higher-Order Chromatin Structures at Different Epigenomic States in Single Mammalian Cells. Cell Rep. 2018, 24 (4), 873–882. https://doi.org/10.1016/j.celrep.2018.06.085.

(46) Cremer, T.; Cremer, M.; Hübner, B.; Strickfaden, H.; Smeets, D.; Popken, J.; Sterr, M.; Markaki, Y.; Rippe, K.; Cremer, C. The 4D Nucleome: Evidence for a Dynamic Nuclear Landscape Based on Co-Aligned Active and Inactive Nuclear Compartments. FEBS Lett. 2015, 589 (20PartA), 2931–2943. https://doi.org/10.1016/j.febslet.2015.05.037.

(47) Li, L.; Lyu, X.; Hou, C.; Takenaka, N.; Nguyen, H. Q.; Ong, C.-T.; Cubeñas-Potts, C.; Hu, M.; Lei, E. P.; Bosco, G.; et al. Widespread Rearrangement of 3D Chromatin Organization Underlies Polycomb-Mediated Stress-Induced Silencing. Mol. Cell 2015, 58 (2), 216–231. https://doi.org/10.1016/j.molcel.2015.02.023.

(48) Corces, M. R.; Granja, J. M.; Shams, S.; Louie, B. H.; Seoane, J. A.; Zhou, W.; Silva, T. C.; Groeneveld, C.; Wong, C. K.; Cho, S. W.; et al. The Chromatin Accessibility Landscape of Primary Human Cancers. Science 2018, 362 (6413), eaav1898. https://doi.org/10.1126/science.aav1898.

(49) Benedetti, F.; Dorier, J.; Burnier, Y.; Stasiak, A. Models That Include Supercoiling of Topological Domains Reproduce Several Known Features of Interphase Chromosomes. Nucleic Acids Res. 2014, 42 (5), 2848–2855. https://doi.org/10.1093/nar/gkt1353.

(50) Bianco, S.; Lupiáñez, D. G.; Chiariello, A. M.; Annunziatella, C.; Kraft, K.; Schöpflin, R.; Wittler, L.; Andrey, G.; Vingron, M.; Pombo, A.; et al. Polymer Physics Predicts the Effects of Structural Variants on Chromatin Architecture. Nat. Genet. 2018, 50 (5), 662–667. https://doi.org/10.1038/s41588-018-0098-8.

(51) Sachs, R. K.; van den Engh, G.; Trask, B.; Yokota, H.; Hearst, J. E. A Random-Walk/Giant-Loop Model for Interphase Chromosomes. Proc. Natl. Acad. Sci. 1995, 92 (7), 2710–2714. https://doi.org/10.1073/pnas.92.7.2710.

(52) Frigola, J.; Song, J.; Stirzaker, C.; Hinshelwood, R. A.; Peinado, M. A.; Clark, S. J. Epigenetic Remodeling in Colorectal Cancer Results in Coordinate Gene Suppression across an Entire Chromosome Band. Nat. Genet. 2006, 38 (5), 540– 549. https://doi.org/10.1038/ng1781.

(53) Jones, P. A.; Baylin, S. B. The Epigenomics of Cancer. Cell 2007, 128 (4), 683–692. https://doi.org/10.1016/j.cell.2007.01.029.

(54) Taberlay, P. C.; Achinger-Kawecka, J.; Lun, A. T. L.; Buske, F. A.; Sabir, K.; Gould, C. M.; Zotenko, E.; Bert, S. A.; Giles, K. A.; Bauer, D. C.; et al. Three-Dimensional Disorganization of the Cancer Genome Occurs Coincident with Long-Range Genetic and Epigenetic Alterations. Genome Res. 2016, 26 (6), 719–731. https://doi.org/10.1101/gr.201517.115.

(55) Wani, A. H.; Boettiger, A. N.; Schorderet, P.; Ergun, A.; Münger, C.; Sadreyev, R. I.; Zhuang, X.; Kingston, R. E.; Francis, N. J. Chromatin Topology Is Coupled to Polycomb Group Protein Subnuclear Organization. Nat. Commun. 2016, 7, 10291. https://doi.org/10.1038/ncomms10291.

(56) Liu, Z.; Tjian, R. Visualizing Transcription Factor Dynamics in Living Cells. J. Cell Biol. 2018, 217 (4), 1181–1191. https://doi.org/10.1083/jcb.201710038.

(57) Ganji, M.; Shaltiel, I. A.; Bisht, S.; Kim, E.; Kalichava, A.; Haering, C. H.; Dekker, C. Real-Time Imaging of DNA Loop Extrusion by Condensin. Science 2018, 360 (6384), 102–105. https://doi.org/10.1126/science.aar7831.

(58) Alipour, E.; Marko, J. F. Self-Organization of Domain Structures by DNA-Loop-Extruding Enzymes. Nucleic Acids Res. 2012, 40 (22), 11202–11212. https://doi.org/10.1093/nar/gks925.

(59) Banigan, E. J.; Mirny, L. A. Limits of Chromosome Compaction by Loop-Extruding Motors: bioRxiv 2018. https://doi.org/10.1101/476424.

(60) Naumova, N.; Imakaev, M.; Fudenberg, G.; Zhan, Y.; Lajoie, B. R.; Mirny, L. A.; Dekker, J. Organization of the Mitotic Chromosome. Science 2013, 342 (6161), 948–953. https://doi.org/10.1126/science.1236083.

(61) Nozaki, T.; Imai, R.; Tanbo, M.; Nagashima, R.; Tamura, S.; Tani, T.; Joti, Y.; Tomita, M.; Hibino, K.; Kanemaki, M. T.; et al. Dynamic Organization of Chromatin Domains Revealed by Super-Resolution Live-Cell Imaging. Mol. Cell 2017, 67 (2), 282–293.e7. https://doi.org/10.1016/j.molcel.2017.06.018.

(62) Wang, K. C.; Yang, Y. W.; Liu, B.; Sanyal, A.; Corces-Zimmerman, R.; Chen, Y.; Lajoie, B. R.; Protacio, A.; Flynn, R. A.; Gupta, R. A.; et al. A Long Noncoding RNA Maintains Active Chromatin to Coordinate Homeotic Gene Expression. Nature 2011, 472 (7341), 120–124. https://doi.org/10.1038/nature09819.

(63) Genereux, J. C.; Boal, A. K.; Barton, J. K. DNA-Mediated Charge Transport in Redox Sensing and Signaling. J. Am. Chem. Soc. 2010, 132 (3), 891–905. https://doi.org/10.1021/ja907669c.

(64) Imai, R.; Nozaki, T.; Tani, T.; Kaizu, K.; Hibino, K.; Ide, S.; Tamura, S.; Takahashi, K.; Shribak, M.; Maeshima, K. Density Imaging of Heterochromatin in Live Cells Using Orientation-Independent-DIC Microscopy. Mol. Biol. Cell 2017, 28 (23), 3349–3359. https://doi.org/10.1091/mbc.e17-06-0359.

(65) Berger, S. L.; Kouzarides, T.; Shiekhattar, R.; Shilatifard, A. An Operational Definition of Epigenetics. Genes Dev. 2009, 23 (7), 781–783. https://doi.org/10.1101/gad.1787609.

(66) Cortini, R.; Barbi, M.; Caré, B. R.; Lavelle, C.; Lesne, A.; Mozziconacci, J.; Victor, J.-M. The Physics of Epigenetics. Rev. Mod. Phys. 2016, 88 (2). https://doi.org/10.1103/RevModPhys.88.025002.

(67) Reinberg, D.; Vales, L. D. Chromatin Domains Rich in Inheritance. Science 2018, 361 (6397), 33–34. https://doi.org/10.1126/science.aat7871.

(68) Yadav, T.; Quivy, J.-P.; Almouzni, G. Chromatin Plasticity: A Versatile Landscape That Underlies Cell Fate and Identity. Science 2018, 361 (6409), 1332–1336. https://doi.org/10.1126/science.aat8950.

(69) Kremer, J. R.; Mastronarde, D. N.; McIntosh, J. R. Computer Visualization of Three-Dimensional Image Data Using IMOD. J. Struct. Biol. 1996, 116 (1), 71–76. https://doi.org/10.1006/jsbi.1996.0013.

(70) Gürsoy, D.; De Carlo, F.; Xiao, X.; Jacobsen, C. TomoPy: A Framework for the Analysis of Synchrotron Tomographic Data. J. Synchrotron Radiat. 2014, 21 (5), 1188–1193. https://doi.org/10.1107/S1600577514013939.

