## Supplemental Information for "Physical and data structure of 3D genome"

### Contact scaling of fractal chain

It has been argued that chromatin is a crumpled globule[1], latter termed fractal (self-similar[2]) globule (FG), so that the long biopolymer could function without entangling and knotting. Computer simulation[3] showed that FG has a contact probability ($P_c$) scaling factor $s$=-1, namely $P_c(L) \propto L^{-1}$, where $L$ is contour distance between two loci on the polymer. We recently based on scale-invariance properties proved that for a generic fractal chain, the $P_c$ scaling factor $s$ is inverse to the fractal dimension $D$ of the chain[4]:

$$s = -\frac{E}{D} \tag{S1}$$

where $E$ is the dimension of space where the fractal chain is embedded. Concerning models of chromatin structure, we have $E$=3 for three-dimensional space. As a special case, FG has a fractal dimension of $D$=3 and therefore $s$=-1 according to our theory, consistent with simulation result reported in literature[3].

To test our scaling theory for more generic cases, we here carried out numerical calculations of the contact probability for random walks (RWs) and Lévy-flights (LFs) in both 2D and 3D space. The random walks have fixed fractal dimension of $D$=2. For LFs we used fractal dimensions 1.2, 1.5 and 1.8, respectively. For all the cases, we generated trajectories of 1 million steps. The RWs have a constant step size, and the LFs have power law distributions of their step size with the exponents determined by their fractal dimensions. We chose the fixed step size of RWs and the minimal step size of LFs to be 1 and employed a contact cutoff to be 3 for the scaling analyses. The asymptotic behaviors of all the cases tested are shown in the colored solid lines in Fig. S1. The dash lines with their exponents predicted by Eq. S1 align well with the numerical results.

Since $s$ and $D$ determine the contact frequency and structural heterogeneity of the fractal chain, respectively, Eq. S1 means that it is impossible for a polymer to simultaneously satisfy (1) high contact frequency, (2) structural heterogeneity and (3) self-similarity. To understand this intuitively: a highly heterogeneous (small $D$) fractal chain such as the LF fills space with self-similar clusters of low inter-cluster contact frequencies[5], whereas a fractal chain of high contact frequencies, such as the FG, tends to occupy the space in a homogeneous way (large $D$) despite its compartmentalization along the polymer contour. Note that there is a subtle yet important difference between compartmentalization and clustering: clusters are spatially isolated compartments. In other words, compartmentalization is a weaker property than clustering, and without isolation it cannot render density heterogeneity in space. The fact that interphase chromatin houses both frequent long-range contacts[6] and heterogeneous local DNA densities[7] thus leads us to conclude that its folding structure is not self-similar. It is also worth noting that since $D \leq E$, we have $s$=-1 as the upper bound for the scaling of fractal polymers. However, at small genomic range, it has been observed[8] that chromatin has an abnormally slow decay of contact probability with $s$>-1, which cannot be explained by fractal chain models.

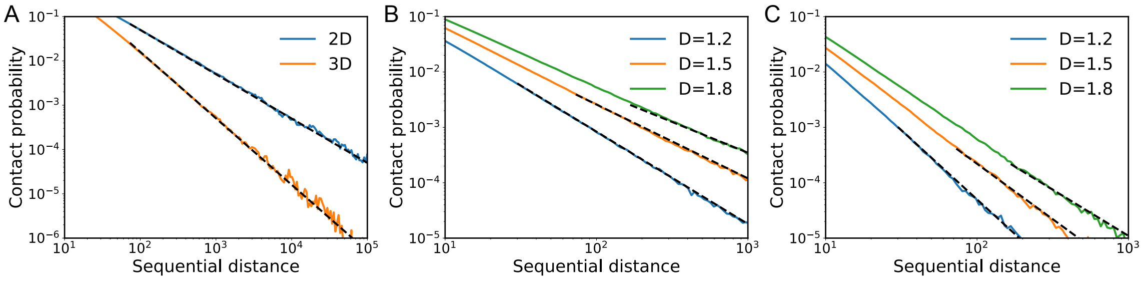


**Figure S1. Contact probabilities of RWs and LFs in 2D and 3D space.** Numerical results are shown in solid lines and theoretical predictions in dash lines (**A**) RWs in 2D and 3D space. (**B**) LFs of varying fractal dimensions in 2D space. (**C**) LFs of varying fractal dimensions in 3D space.

### Tree domains and TADs

A typical 3D structure of a 3Mb segment of SRRW (1500 steps) is shown in Fig. S2A. The local segment is clearly folded into a bundle of clusters at nano-scale. Allowing such conformation to relax with preserved topology offers a conformational set (1000 conformations) on which we can obtain a population-level contact map featuring TAD-like domains as shown in Fig. S2B. Such 2D contact patterns are the statistical consequence of hierarchical 3D structures linked and isolated by stretched DNA. In this example, the 3Mb contact map divides itself into two large domains isolated from each other by extended backbone segments. Inside these large domains, similar genomic isolations recur at smaller scales, leading to a hierarchical organization of domains. From a loop point of view, this means that large loops enclose small loops in a hierarchical manner. Such tree-like nested loops avoid entangling and knotting of long DNA in a crowded and disordered environment. However, tree domains are not simply nested loops as they are rich in chromatin nanoclusters or clutches. Our model suggests that distal chromatin nanoclusters can undergo long-range higher-order interactions to form tree nodes that stabilize the topological domains in single cells. The population-average TADs have been explained by the loop extrusion hypothesis[10], in which large loops are formed by extrusion machinery (likely realized by CTCF and cohesin). However, at the single-cell level, loop extrusion does not produce clusters that are found to be ubiquitous under super-resolution microscopy even with cohesin knocked out[11]. In contrast, SRRW naturally assumes a clustered morphology at single-cell level.

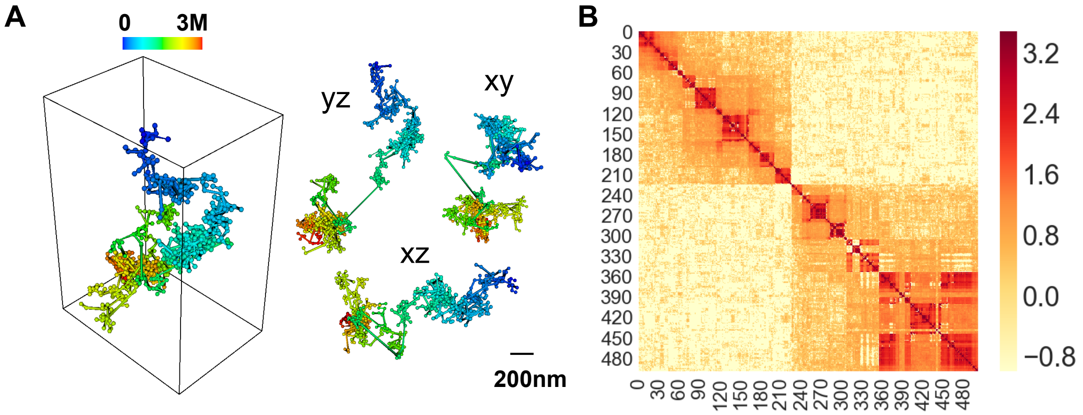


**Figure S2. Structure of local SRRW segment and ensemble-averaged contact map.** (**A**) 3D structure of a 3Mb segment of SRRW and its xyz projections. (**B**) Contact map of the 3Mb SRRW segment over 1000 random conformations with preserved topology. Color bar in logarithm scale.

It is worth noting that tree domains are similar but not equivalent to Hi-C TADs: while Hi-C TADs are contact patterns that emerge over the sampling of millions of cells, trees domains predicted by our model are complexes of loops and clusters at the single-cell level. Due to the loss of information from 3D structures to 2D contact maps, qualitatively different 3D structural predictions can result in similar 2D contact patterns. Since Hi-C is sensitive to both topological contacts (contacts underpinning topological constraints) and natural contacts (transient contacts with less functional significance), it is hard to identify specific folding modes based on only ensemble-averaged contact maps. For example, a 1Mb-level TAD pattern could be possibly resulted from either a large tree-like topological domain or from a small non-topological compartment. Therefore, it is important for modeling efforts to account for both Hi-C and imaging observations.

Recent experimental evidence shows that contact loops are dynamic[9], meaning they can form and break, allowing rapid epigenetic reconfiguration. Given this dynamic picture, we posit that, like chemical reaction, small tree domains can group into large tree domains, which can in reverse ungroup into small ones. The contact map in Fig. S2B is based on a frozen topology without any regrouping of tree domains and therefore could have overestimated the isolation across domains, which renders an over-hierarchical contact map at population-level. Nevertheless, as a proof of concept, it clearly demonstrated how single-cell level tree domains could lead to TAD patterns at population-level.

**Non-Gaussian statistics**

Heterogeneity and non-Gaussian statistics are hallmarks of living systems. The intrinsic heterogeneity of SRRW is tightly associated with its non-Gaussian statistics. For example, the end-to-end distance ($R$) distribution of a sample DNA segment of 100kb predicted by SRRW is significantly deviated from normal, as shown in green in Fig. S3, with high probability of small $R$ due to self-returning and a heavy tail of stretched DNA with large $R$. Compared to the counterpart from a RW, the 100kb segment of SRRW is much shorter on average, signifying strong compaction. The existence of the long tail allows the compacted domains to be spatially separated.

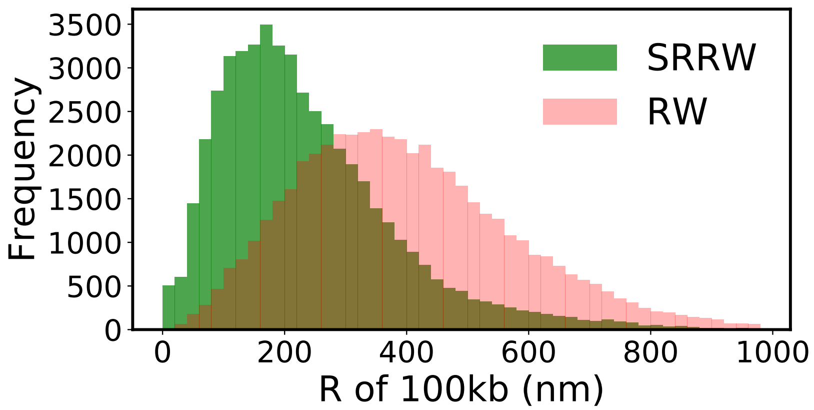


**Figure S3. End-to-end distance distributions of 100kb SRRW segment (green) and of 100kb RW segment (red).**

### Effect of $\alpha$ on the statistics of tree domains

In the SRRW model, $\alpha$ governs the global folding architecture of the modeled chromatin and impacts the heterogeneity of the local DNA packing, which has been demonstrated and discussed in the main text. Here we show more numerical results on the effect of $\alpha$. As shown in Fig. S4, the distribution of the physical size of tree domains and the distribution of the genomic size of the tree nodes are both sensitive to $\alpha$. In Fig. S4A, the physical size of the tree domain is characterized by the radius of gyration ($R_g$) and normalized by the genomic size of the domain. Larger tree domains are more hierarchical in structure and contain more high-order contacts.

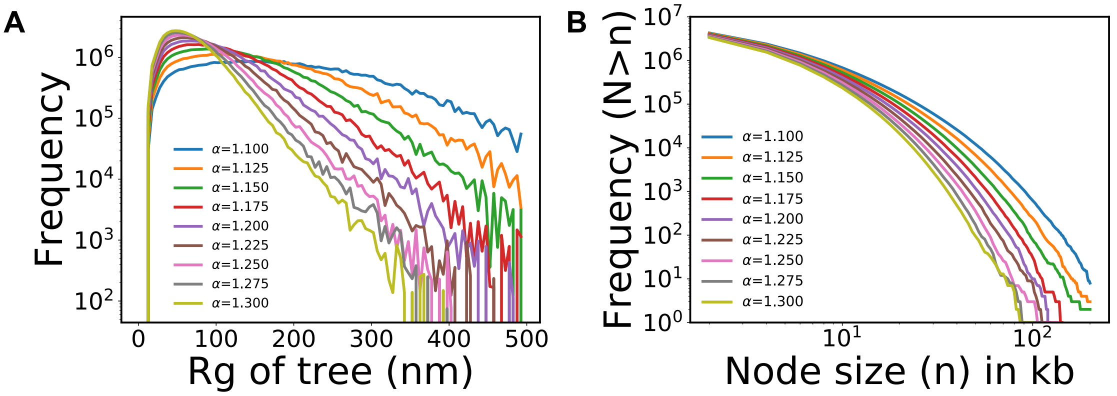


**Figure S4. Effect of $\alpha$ on the tree domains.** (**A**) $R_g$ distribution of tree domains normalized by their genomic sizes. (**B**) Genomic size distribution of the tree nodes.

As discussed in the main text, there is an anti-correlation between the effective mass scaling $D$ and the contact scaling $s$. Fig. S5 shows the data sets from which we extracted $D$ and $s$.

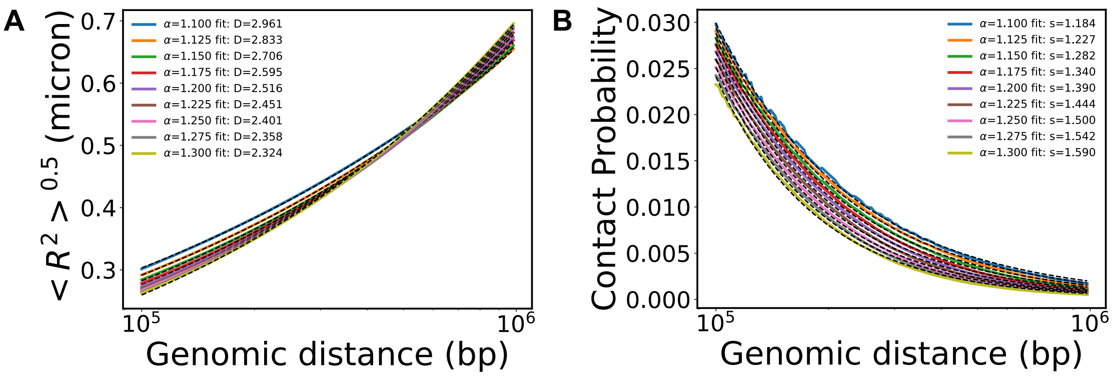

**Figure S5. $D$ and $s$ from the modeled chromatin at varying $\alpha$.** (**A**) End-to-end distance scalings and their power-law fittings from 100kb to 1000kb. (**B**) Contact probability scalings and fittings for the same genomic range.

### ChromSTEM materials and methods

1. Cell culture and EM sample preparation.

Human lung carcinoma cells (A549, ATCC ® CCL-185 ™) and Human skin fibroblast cells (BJ, ATCC CRL®-2522™) were cultured on 35 mm MatTek dishes (MatTek Corp) in xxx in 5% $CO_2$ and at physiological oxygen level. Cells were rinsed in Hanks's balanced salt solution without calcium and mageneisum then fixed with 2.5% EM grade glutaraldehyde (EMS) in 5mM $CaCl_2$, 0.1M sodium cacodylate buffer, pH 7.4 for 5 min at room temperature. Then fresh fixative was used to continue fixing the cells on ice for an hour. From this step, the cells were kept cold on ice or on a cold stage with temperature ranges from 4 $^{o}$C to 10 $^{o}$C. The ChromEM staining and embedding follows published protocol[7]. 100 nm A549 cell sections and 50 nm BJ cell sections were made by ultramicrotomy (Leica, UC7) and mounted on copper mesh grids with Formvar/carbon film (EMS).

2. EM imaging and tomography reconstruction

Solution of 10 nm colloidal gold fiducial markers was deposited onto both slide of the sample prior to imaging. A 200 kV STEM (HD2300, HITACHI) was employed to collect the dual-tilt series in high angle annular dark field (HAADF) mode. During each tilt series, the sample was tilted from -60$^{o}$ to 60$^{o}$ with a step size of 2$^{o}$. Between the two-tilt series, the sample was taken out of the microscope and rotated around 90$^{o}$ manually. A low dose imaging condition was conducted to minimize beam damage. The tilt series were aligned in IMOD[12] using fiducial markers. After alignment, each tilt series were reconstructed separately using penalized maximum likelihood algorithm in Tomopy[13]. IMOD was used again to combine the two reconstructed tomograms to minimize artifacts from missing wedge. The BJ cell sections were imaged by a TEM (HT7700, HITACHI) operated at 80 kV. The brightfield micrographs were recorded with pixel resolution of 5 nm for the nucleus region for 20 cells.

3. Chromatin volume concentration (CVC) calculation and nanocluster size analysis

The automated chromatin segmentation was conducted using CLAHE (100 nm) – Li in FIJI as previously described. We used the same definition of the chromatin volume concentration (CVC) as Ou's work [1] but with a moving window summation with a three-dimension window size of 99 nm and a stride of 3 nm. From the CVC map, we observed nanoclusters with similar values in CVC, and measured the full width half maximum (FWHM) of the line profile of CVC as the cluster diameter. 29 clusters were selected, and the histogram of the cluster size was plotted.

4. Mass scaling analysis with TEM images

The 2D-mass scaling was calculated for each tomogram slice. Starting with a random position around the center of chromatin and draw a circle with radius r, the total chromatin (M) encapsulated by the circle can be calculated. The mass-scaling is the relation between the radius r and the total chromatin (M) encapsulated. The mass scaling M(r) follows a power law and for fractal structure in specific, the power is smaller than 2 for the 2D slice and smaller than 3 for the 3D case.

**ChromSTEM images**

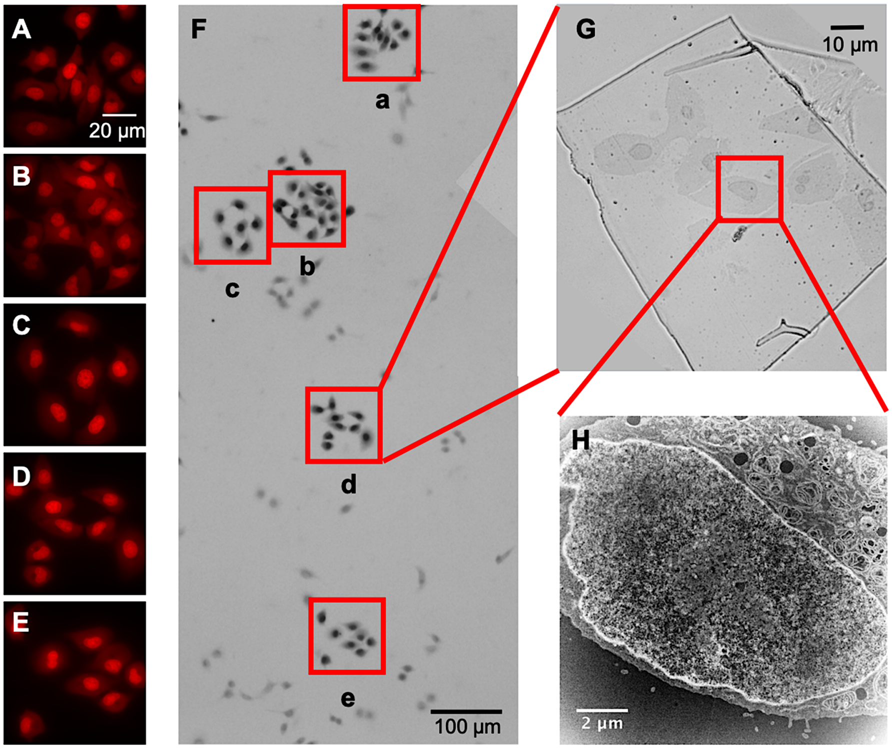


**Figure S6. ChromEM sample preparation and STEM imaging on A540 cells.** (**A**) to (**E**), fluorescence image of A549 cells with DRAQ5 label during photo-bleaching. (**F**) Optical bright field image of A549 cells after resin embedding. Spots after photo-bleaching (a to e) showed darker contrast compare to neighboring cells without photo-bleaching. (**G**) Optical bright filed image of one thin section of A549 cells prepared by ChromEM protocol. (**H**) STEM HAADF imaging of 100 nm ultra-thin resin section of one A549 cell.

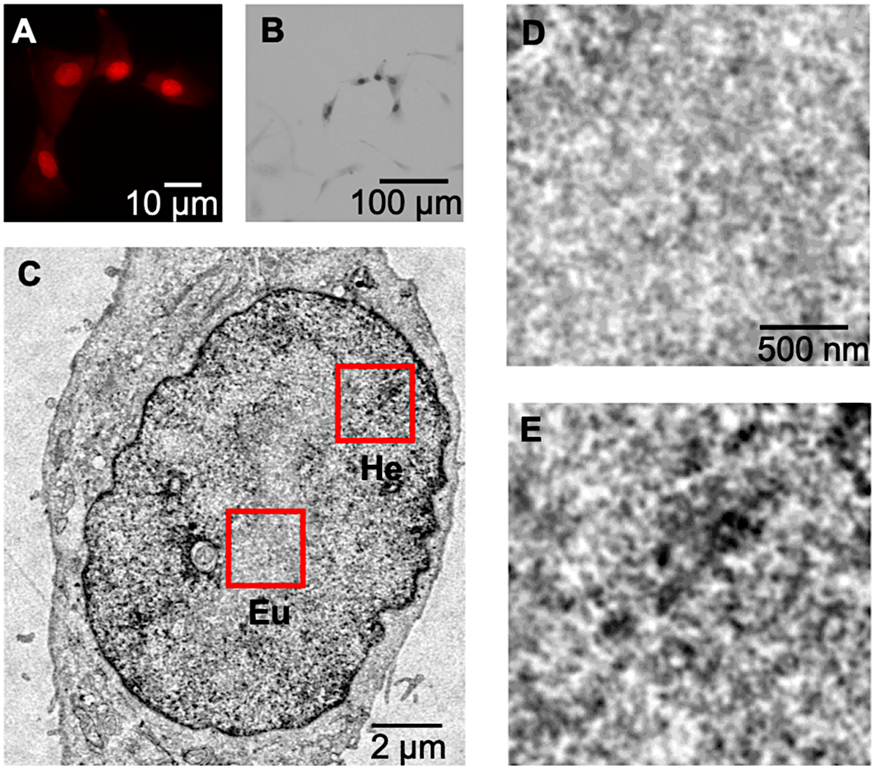


**Figure S7. ChromEM sample preparation and TEM imaging on BJ cells.** (**A**) Fluorescence image of BJ cells labeled with DRAQ5 during photo-bleaching. (**B**) Optical bright field image after resin embedding for corresponding cells in (A). Compared to neighboring cells, the cells with photo-bleaching showed significantly darker contrast. (**C**) TEM image of a BJ cell with chromEM preparation. Euchromatin (Eu) and heterochromatin (He) were identified based on the DNA density. (**D**) Magnetified image of the euchromatin (red square, Eu) in (C). (**E**) Magnified image of the heterochromatin (red square, He) in (C). The euchromatin showed lighter contrast compared to the heterochromatin.

### PWS materials and methods

HeLa cells (Figs. 5d, S8) were grown on 35 mm glass bottom dishes until approximately 70% confluent. Cells were imaged with Partial Wave Spectroscopic (PWS) microscopy while in a stage-top incubator chamber (DMIW chamber, Tokai Hit) which regulated temperature (37° C) and CO2 (5%). PWS has been described in detail in previous works[14–17]. Briefly, PWS microscopy allows interrogation of macromolecular organization in both fixed and living cells[13–15] with length-scale sensitivity in the range of 20–200 nm. Light from a broadband source (X-Cite

120LED) is coupled to a commercial microscope base (Eclipse Ti-U with perfect-focus system, Nikon or DMIRB, Leica) and is focused onto the sample with a small illumination numerical aperture (~0.5). The backscattered light is collected with a high NA objective (1.49 or 1.4) and sent through a spectral filter (CRi VariSpec LCTF) and imaged with a sCMOS or EMCCD camera (ORCA-Flash4.0, Hamamatsu or ImageEM C9000-13, Hamamatsu). The scaling of chromatin packing (*D*) is calculated from variances in the spectrally resolved backscattered light[14–16]. For the heat shock experiments, HCT-116 cells (Figs. 6L, M) were grown to confluence in McCoy's 5A medium (Lonza) supplemented with 10% FBS and 1X Penicillin-Streptomycin at 37 °C. Cells from passage 5-20 were used for the experiment. Cells incubated at 37 °C (with 5% $CO_2$) were treated as controls. For heat shock, each dish was imaged using PWS microscopy, incubated at 42 °C (with 5% $CO_2$) for 1 hour and then immediately imaged afterwards. All the experiments were blinded.

### PWS images

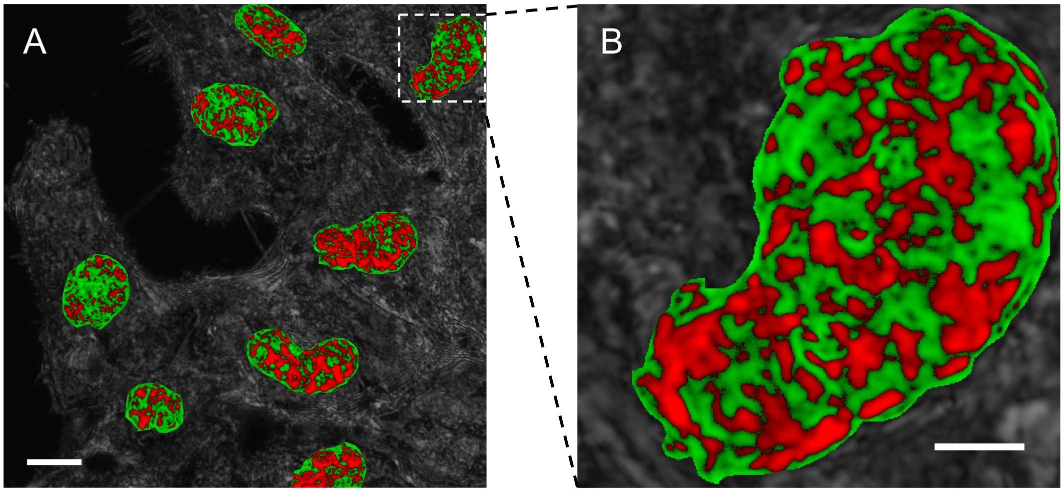


**Figure S8. Live-cell PWS image of HeLa cells.** (**A**) Multi-cell image. The scaling of chromatin packing within the nuclei is shown in red and green color. Regions colored in green have lower scaling of chromatin packing while regions in red are packing domains with higher scaling of chromatin packing. Scale bar: 10 μm. (**B**) A zoomed in image from A showing the distribution of packing domains within a single nucleus. Scale bar: 3 μm.

### Heat shock Hi-C analysis

Hi-C data was taken from the publicly available Hi-C data[20] on Kc167 *Drosophila* cells (GEO access code GSE63518) grown at normal temperature (25°C) and exposed to heat shock (36.5°C) for 20 min. Hi-C data from 4 control and 2 heat shock biological replicates were averaged and the intra-chromosomal contact probability scaling (s) with increasing genomic distance was quantified for the entire genome with a resolution of 5 kb. Hi-C contact maps for chromosome (chr) 2L were produced by combining contacts from biological replicates of control and heat shock data and were plotted as heat maps. The Hi-C heatmaps were plotted at multiple resolutions: single-fragment (3.5 kb) resolution for a 2 Mb section of chr 2L and 7 kb resolution for the entire chromosome.

### References


(1) Grosberg, A.; Rabin, Y.; Havlin, S.; Neer, A. Crumpled Globule Model of the Three-Dimensional Structure of DNA. *Europhys. Lett. EPL* **1993**, *23* (5), 373–378. https://doi.org/10.1209/0295-5075/23/5/012.

(2) Mandelbrot, B. B. *The Fractal Geometry of Nature*; W.H. Freeman: San Francisco, 1982.

(3) Mirny, L. A. The Fractal Globule as a Model of Chromatin Architecture in the Cell. *Chromosome Res.* **2011**, *19* (1), 37–51. https://doi.org/10.1007/s10577-010-9177-0.

(4) Huang, K.; Backman, V.; Szleifer, I. Interphase Chromatin as a Self-Returning Random Walk: Can DNA Fold into Liquid Trees? **2018**. https://doi.org/10.1101/413872.

(5) *Lévy Flights and Related Topics in Physics*; Shlesinger, M. F., Zaslavsky, G. M., Frisch, U., Eds.; Araki, H., Brézin, E., Ehlers, J., Frisch, U., Hepp, K., Jaffe, R. L., Kippenhahn, R., Weidenmüller, H. A., Wess, J., Zittartz, J., et al., Series Eds.; Lecture Notes in Physics; Springer Berlin Heidelberg: Berlin, Heidelberg, 1995; Vol. 450. https://doi.org/10.1007/3-540-59222-9.

(6) Rao, S. S. P.; Huntley, M. H.; Durand, N. C.; Stamenova, E. K.; Bochkov, I. D.; Robinson, J. T.; Sanborn, A. L.; Machol, I.; Omer, A. D.; Lander, E. S.; et al. A 3D Map of the Human Genome at Kilobase Resolution Reveals Principles of Chromatin Looping. *Cell* **2014**, *159* (7), 1665–1680. https://doi.org/10.1016/j.cell.2014.11.021.

(7) Ou, H. D.; Phan, S.; Deerinck, T. J.; Thor, A.; Ellisman, M. H.; O'Shea, C. C. ChromEMT: Visualizing 3D Chromatin Structure and Compaction in Interphase and Mitotic Cells. *Science* **2017**, *357* (6349), eaag0025. https://doi.org/10.1126/science.aag0025.

(8) Sanborn, A. L.; Rao, S. S. P.; Huang, S.-C.; Durand, N. C.; Huntley, M. H.; Jewett, A. I.; Bochkov, I. D.; Chinnappan, D.; Cutkosky, A.; Li, J.; et al. Chromatin Extrusion Explains Key Features of Loop and Domain Formation in Wild-Type and Engineered Genomes. *Proc. Natl. Acad. Sci.* **2015**, *112* (47), E6456–E6465. https://doi.org/10.1073/pnas.1518552112.

(9) Hansen, A. S.; Cattoglio, C.; Darzacq, X.; Tjian, R. Recent Evidence That TADs and Chromatin Loops Are Dynamic Structures. *Nucleus* **2018**, *9* (1), 20–32. https://doi.org/10.1080/19491034.2017.1389365.

(10) Fudenberg, G.; Imakaev, M.; Lu, C.; Goloborodko, A.; Abdennur, N.; Mirny, L. A. Formation of Chromosomal Domains by Loop Extrusion. *Cell Rep.* **2016**, *15* (9), 2038–2049. https://doi.org/10.1016/j.celrep.2016.04.085.

(11) Bintu, B.; Mateo, L. J.; Su, J.-H.; Sinnott-Armstrong, N. A.; Parker, M.; Kinrot, S.; Yamaya, K.; Boettiger, A. N.; Zhuang, X. Super-Resolution Chromatin Tracing Reveals Domains and Cooperative Interactions in Single Cells. *Science* **2018**, *362* (6413), eaau1783. https://doi.org/10.1126/science.aau1783.

(12) Kremer, J. R.; Mastronarde, D. N.; McIntosh, J. R. Computer Visualization of Three-Dimensional Image Data Using IMOD. *J. Struct. Biol.* **1996**, *116* (1), 71–76. https://doi.org/10.1006/jsbi.1996.0013.

(13) Gürsoy, D.; De Carlo, F.; Xiao, X.; Jacobsen, C. TomoPy: A Framework for the Analysis of Synchrotron Tomographic Data. *J. Synchrotron Radiat.* **2014**, *21* (5), 1188–1193. https://doi.org/10.1107/S1600577514013939.
(14) Cherkezyan, L.; Capoglu, I.; Subramanian, H.; Rogers, J. D.; Damania, D.; Taflove, A.; Backman, V. Interferometric Spectroscopy of Scattered Light Can Quantify the Statistics of Subdiffractional Refractive-Index Fluctuations. *Phys. Rev. Lett.* **2013**, *111* (3). https://doi.org/10.1103/PhysRevLett.111.033903.
(15) Cherkezyan, L.; Subramanian, H.; Backman, V. What Structural Length Scales Can Be Detected by the Spectral Variance of a Microscope Image? *Opt. Lett.* **2014**, *39* (15), 4290. https://doi.org/10.1364/OL.39.004290.
(16) Chandler, J. E.; Cherkezyan, L.; Subramanian, H.; Backman, V. Nanoscale Refractive Index Fluctuations Detected via Sparse Spectral Microscopy. *Biomed. Opt. Express* **2016**, *7* (3), 883. https://doi.org/10.1364/BOE.7.000883.
(17) Chandler, J. E.; Stypula-Cyrus, Y.; Almassalha, L.; Bauer, G.; Bowen, L.; Subramanian, H.; Szleifer, I.; Backman, V. Colocalization of Cellular Nanostructure Using Confocal Fluorescence and Partial Wave Spectroscopy. *J. Biophotonics* **2017**, *10* (3), 377–384. https://doi.org/10.1002/jbio.201500298.
(18) Almassalha, L. M.; Bauer, G. M.; Chandler, J. E.; Gladstein, S.; Cherkezyan, L.; Stypula-Cyrus, Y.; Weinberg, S.; Zhang, D.; Thusgaard Ruhoff, P.; Roy, H. K.; et al. Label-Free Imaging of the Native, Living Cellular Nanoarchitecture Using Partial-Wave Spectroscopic Microscopy. *Proc. Natl. Acad. Sci.* **2016**, *113* (42), E6372–E6381. https://doi.org/10.1073/pnas.1608198113.
(19) Zhou, X.; Almassalha, L.; Li, Y.; Eshein, A.; Cherkezyan, L.; Viswanathan, P.; Subramanian, H.; Szleifer, I.; Backman, V. Preservation of Cellular Nano-Architecture by the Process of Chemical Fixation for Nanopathology. *bioRxiv* **2018**. https://doi.org/10.1101/371286.
(20) Li, L.; Lyu, X.; Hou, C.; Takenaka, N.; Nguyen, H. Q.; Ong, C.-T.; Cubeñas-Potts, C.; Hu, M.; Lei, E. P.; Bosco, G.; et al. Widespread Rearrangement of 3D Chromatin Organization Underlies Polycomb-Mediated Stress-Induced Silencing. *Mol. Cell* **2015**, *58* (2), 216–231. https://doi.org/10.1016/j.molcel.2015.02.023.
